# Coupling of Coastal Activity with Tidal Cycles is Stronger in Tool-using Capuchins (*Cebus capucinus imitator*)

**DOI:** 10.1101/2022.12.22.521421

**Authors:** Zoë Goldsborough, Margaret C. Crofoot, Shauhin E. Alavi, Sylvia F. Garza, Evelyn Del Rosario-Vargas, Kate Tiedeman, Claudio M. Monteza-Moreno, Brendan J. Barrett

## Abstract

Terrestrial mammals exploiting coastal resources must cope with the challenge that resource availability and accessibility fluctuate with tidal cycles. Tool use can improve foraging efficiency and provide access to structurally protected resources that are otherwise unavailable (e.g., mollusks and fruits). To understand how variable accessibility of valuable resources shapes behavioral patterns, and whether tool use aids in the efficient exploitation of intertidal resources, we compared the relationship between tidal cycles and activity patterns of tool-using vs. non-tool-using groups of white-faced capuchin monkeys on Jicarón Island in Coiba National Park, Panama. Although only a single group of capuchins on Jicarón uses tools, all coastal groups forage on intertidal resources. Using data from >3 years of camera trapping at varying distances from the coast, we found that capuchins on Jicarón showed increased coastal activity during specific parts of the tidal cycle, and that this relationship differed between tool-using and non-tool-using groups, as well as between seasons. Activity patterns of tool-using capuchins were more strongly and consistently tied to tidal cycles compared to non-tool-users, indicating that tool use might allow for more efficient exploitation of tidal resources. Our findings highlight the potential of tool use to aid niche expansion.

## Introduction

Animals that are dietary generalists are buffered against the negative consequences of environmental change and fluctuating resource availability (Hill & Winder, 2019). Dietary generalists have been shown to be more innovative and better at spatial reasoning than specialist congeners helping them to access patchy resources (Henke-von der Malsburg et al., 2020, 2021), and have also fared better than specialists in the face of anthropogenic change (Clavel et al., 2011). Maritime mammals (*sensu* Carlton & Hodder, 2003) are one example of dietary generalists: these are populations of foragers who exploit resources in the intertidal zones, despite intertidal areas being an unlikely selective pressure in their evolutionary past, and who typically also forage in other habitats.

While all animals must deal with spatiotemporal variability in choosing when and where to forage, terrestrial foragers exploiting intertidal resources face additional challenges. Tidal cycles are asynchronously cyclical: tidal peaks occur predictably, but the timing of low and high tides shifts daily and the magnitude of tidal change is affected by a myriad of interacting processes. Resource accessibility also varies depending on season and weather. Different prey species may show intra-annual variability in abundance due to migration or breeding patterns (Collin & Ochoa, 2016; Gyory, 2010). Prey accessibility can be limited by sunlight and temperature, which can heat substrates to > 45 °C and impact tidal pool salinity causing prey to conceal themselves or move towards the ocean (Abele, 1974; Garrity, 1984; Trowbridge, 1994). Further, the intertidal zone in tropical regions is often exposed and hot, which potentially puts individuals with reduced adaptation to cope with hot substrates, at risk of thermal stress when traversing the intertidal zone. Although a taxonomically diverse set of mammalian taxa are known to exploit intertidal resources (e.g. primates [Gumert & Malaivijitnond, 2012; Lewis & O’Riain, 2017], carnivores [Simmons et al., 2014; Stander, 2019; Suraci et al., 2017] and rodents [Navarrete & Castilla, 1993]), it is unclear *how* this variable accessibility of valuable resources affects the daily activity patterns of these maritime mammals.

### Tool use improves efficiency of intertidal foraging

The importance of intertidal resources and the extent to which maritime mammals alter their activity patterns to tidal cycles may depend on their ability to exploit the resources in these dynamic habitats. Intertidal resources, such as marine invertebrates and fruits with ocean dispersed seeds, are physically protected and therefore require time-consuming processing (e.g., bivalves, crabs). One way in which maritime mammals could exploit intertidal resources more efficiently and access resources that are otherwise inaccessible to them is by using tools. Tool use facilitates more efficient resource consumption through faster processing, and provides access to structurally protected foods (Biro et al., 2013), and as such allows animals to expand their effective environment. Rocky intertidal areas exposed by low tides provide a wealth of anvil and hammerstone material, in the form of bedrock, driftwood and smooth stones.

A behavior is considered tool use when an animal manipulates an object that is not part of their own body in order to reach a useful outcome (Beck, 1980; Shumaker et al., 2011). Examples of animal tool use in foraging contexts are widespread, ranging from chimpanzees (*Pan troglodytes*) using sticks for termite fishing (Goodall, 1964), to dolphins (*Tursiops sp.*) protecting their rostra with sponges while foraging on the seafloor (Krutzen et al., 2005) and New-Caledonian crows (*Corvus moneduloides)* crafting and using hooked as well as unhooked twig tools to forage on embedded insects (Rutz & St Clair, 2012).

### Maritime mammals face challenges in timing foraging with tidal cycles

How intertidal foraging impacts a maritime mammal’s behavior depends on the importance of these resources in their diets. Carlton & Hodder (2003) distinguish between opportunistic and obligate foraging in the intertidal zone. Opportunistic omnivores can exploit the intertidal zone whenever it is advantageous to them. In contrast, obligate reliance on intertidal resources can be a result of seasonal or systematic impoverishment of terrestrial resources which drives maritime mammals to intertidal resources as an essential addition to their diet (Carlton & Hodder, 2003), making intertidal resources a type of fallback food (Marshall et al., 2009; Marshall & Wrangham, 2007). Opportunistic exploitation of the intertidal zone has different effects on maritime mammals’ activity patterns. An example of opportunistic maritime mammals are Japanese macaques (*Macaca fuscata*), who feed on seafood and only show intertidal foraging linked to tidal cycles in months when terrestrial resources are scarce (Tsuji & Kazahari, 2019). In contrast, black legged kittiwakes (*Rissa tridactyla*) – who are more obligate maritime mammals as they systematically rely on intertidal resources – choose to forage in the intertidal zone and leave their chicks unattended when low tide overlaps with their nesting period (Irons, 1998). Furthermore, island-living arctic foxes (*Alopex lagopus*), whose diet is dominated by fish caught in tidal pools, shape their daily activity pattern around the tides: they forage at low tide and sleep at high tide (Nielsen, 1991). Animals that exploit resources in the intertidal zone may thus benefit from adjusting their daily activity to the tidal cycles.

### The role of tool use in intertidal exploitation

Tool use might aid intertidal exploitation by maritime mammals, as many intertidal resources are structurally protected (e.g., snails, bivalves, coconuts). An example of animals using tools to forage in intertidal zones with greater efficiency are bearded capuchins (*Sapajus apella* and *Sapajus libidinosus*) in mangroves. These capuchins exploit intertidal resources, including crabs and snails, with and without tools (Dos Santos et al., 2019). Notably, while crabs can be eaten without using tools, it appears that snails can only be consumed by tool-using capuchins. Island-living Burmese macaques (*Macaca fascicularis*) exploit tidal resources, such as oysters, by using stone tools and forage in the intertidal zone at higher frequencies during low tide (Gumert & Malaivijitnond, 2012). Intertidal resources may be of differing importance to tool-using and non-tool-using maritime mammals, as tool use can provide access to novel foraging niches (Krützen et al., 2014) or otherwise inaccessible items (Sanz & Morgan, 2013). Thus, tool-using maritime mammals may access more marine prey items than non-tool-using mammals. Alternatively, tool-users might have *less* need for intertidal resources than non-tool-users as they can access more nutrient-rich foods inland as well. In this case, non-tool-using maritime mammals might exploit intertidal resources to a greater extent as they have fewer terrestrial resource options.

### Coiban capuchins are tool-using and non-tool-using maritime mammals

White-faced capuchins in Coiba National Park, Panama (hereafter Coiban capuchins), have been reported to habitually conduct first-order (e.g., pounding of coconuts on anvils; Méndez-Carvajal & Valdés-Díaz, 2017) and second-order tool use (e.g., hammerstone and anvil stone tool use; Barrett et al., 2018; Monteza-Moreno, Dogandžić, et al., 2020). This habitual tool use behavior provides the opportunity to explore: *i)* if capuchins adjust their patterns of activity to coincide with the tides and *ii)* whether this relationship differs between tool-using and non-tool-using capuchins. White-faced capuchins are exploratory dietary generalists who rely heavily on extractive foraging to access structurally protected resources in varied neotropical forests (Moynihan, 1976; Oppenheimer, 1968; Perry & Ordonez Jiménez, 2006). Many of these extractive foraging behaviors are socially-learned (Barrett et al., 2017; Panger et al., 2002; Perry, 2011). Despite decades of studies across multiple field sites (Fedigan & Jack, 2012; Perry et al., 2012), habitual stone-tool use by white-faced capuchins has only been documented in one group of capuchins on the island of Coiba (Monteza-Moreno, Dogandžić, et al., 2020) and one group on the island of Jicarón (Barrett et al., 2018) in Coiba National Park, located ∼30 km off the Pacific coast of Panama. Recently, stone tool use was also described in an urban population of *Cebus albifrons* in Ecuador (Araujo et al., 2021). Coiban capuchins use hammerstone and anvil tool use to access a variety of food items, including sea almonds (*Terminalia catappa*), coconuts (*Cocos nucifera*), Halloween crabs (*Gecarcinus quadratus*), palm fruits (*Bactris major* & *Astrocaryum spp.*), hermit crabs (*Coenobita compressus*), nerite snails (*Nerita sp.)*, and other freshwater mollusks. Coiban capuchins differ from well-studied mainland populations in their degree of terrestriality, likely due to the lack of mammalian predators on the island (Monteza-Moreno, Crofoot, et al., 2020) and live at high population densities (Barrett et al., 2018; Milton & Mittermeier, 1977). Additionally, both Coiba and Jicarón have lower plant richness than similar mainland ecosystems (Ibáñez, 2011; Pérez et al., 1996). On Jicarón, this tool use tradition is highly localized along the coast and limited to one social group (Barrett et al., 2018), despite similar ecological circumstances and no physical barriers between the tool-using capuchin group and other groups of non-tool-using capuchins on the island. Further, both tool-using and non-tool-using capuchins on Jicarón are maritime mammals– that forage on (structurally protected) marine prey items along the coast and in the intertidal zone, such as crabs and snails (Barrett et al., 2018; Video 1). Coiba National Park experiences mixed tides; two low tides of different heights occur every ∼12.5 hours. Thus, the timing of the low tide(s) experienced by capuchins shifts approximately 30 minutes each day. (see Supplementary Figure S1 for the timing of low tides for one month). As such, visits to the intertidal zone at a repeatable time each day is insufficient for maritime mammals to regularly exploit coastal resources.

Here, we investigate how the pattern of Coiban capuchins’ activity varies with shifting tidal cycles, by using camera trap data collected from March 2017-March 2019. We compare how the patterns, relative to distance from coast and tidal cycles, differ between tool-users and non-tool users. As exploitation of intertidal resources likely has a seasonal component (e.g., red deer, Conradt, 2000; and baboons, Lewis & O’Riain, 2017), we examine differences in tidal patterns for tool-using and non-tool-using capuchins between the wet and the dry season. Lastly, capuchin activity at the coast can be affected by diurnal activity and space use patterns, which are likely shaped by temperature and sleeping site location. To gain a more complete picture of capuchins’ activity at the coast, we also consider diurnal capuchin activity (i.e., are capuchins more likely to be at the coast at a specific time of day). By jointly examining these questions in a system with tool-using and non-tool-using capuchins, we provide a first exploration of the relationship between tool use and tidal cycles for a non-human animal in a coastal habitat. This provides insights into how an ecological generalist copes with complicated cycles of resource availability in a non-typical habitat, and provides a much needed comparative study to help understand potential hominin behavior in coastal environments where tool use is unlikely to be preserved (de Chevalier et al., 2022; Marean, 2014).

** Video 1 ** (https://youtu.be/Qwued08S3Xs)

**Video 1:** *Examples of coastal activity by tool-using Coiban capuchins*. Two videos from camera traps at the edge of the coastal vegetation line, showing a) an adult female consuming a Halloween crab in several juveniles traveling on the beach, and b) both foraging on the beach and tool use on a driftwood anvil.

## Methods

### Subjects & Site

Coiba National Park consists of nine islands and over 100 islets located off the Pacific coast of Veraguas Province, Panama. It is a designated UNESCO World Heritage site with endemic animal and plant species (Ibáñez et al., 1997). White-faced capuchins (*Cebus capucinus imitator*) live on the islands of Coiba (50 314 ha), Jicarón (2 002 ha), and Rancheriá (125 ha). These islands are estimated to have been geographically isolated from mainland Panama for 14 000-18 000 years (Titcomb & O’Dea, 2020). The terrestrial mammalian communities in Coiba National Park are depauperate in comparison to forests on the mainland, and mammalian predators are entirely absent. Coiba and Jicarón were used as a penal colony from 1919 until 2004, prior to which the islands were inhabited by indigenous people from 250 CE until about the 16^th^ century (Isaza & Vrba, 2010). In recent years, only the island of Coiba and Rancheriá see constant human occupation at two research stations and a police station; the other islands (including Jicarón) are largely undisturbed (Barrett et al., 2018). Average annual temperature in Coiba National Park is around 26 C°, and rainfall strongly varies seasonally: in the dry season (mid-December to mid-April) there is less than 60 mm of precipitation, while in the wet season there is over 3000 mm of precipitation (Cardiel et al., 1997). Since the start of data collection on Jicarón in 2017, tool use has only been observed to occur along a ∼1 km stretch of coast, likely occupied by a single group of capuchins as documented by camera trap data. Extensive camera trapping and surveys of the coast and in riparian areas yielded no evidence of tool use outside of this stretch of coast. Therefore, we refer to the other surveyed groups on the island as non-tool-using groups.

### Data Collection & Processing

#### Data collection

We analyzed images and videos collected using unbaited camera traps placed at targeted sites (e.g., on tool use anvils or in sea almond groves) on the island of Jicarón between 25 March 2017 and 12 August 2019 in 7 deployments of about ∼4 months (3-5 months). Both still (Reconyx Hyperfire HC600 & HF2X) and video (Reconyx Ultrafire XR6 & XP9) camera traps were used. Camera traps are not fully non-invasive, as they are visually incongruent with the surroundings, and produce sounds or light which may disturb animals (and some individuals/species more than others) (Caravaggi et al., 2020). We purchased infrared rather than white flash camera traps to minimize disturbance to the animals.

Still-image camera traps recorded 10 images per trigger event without any between-trigger delays (∼1 sec between images). Video cameras recorded over a 24-hour period and captured one image and a dynamic video per trigger, which means that the camera trap stops recording after 3 seconds of inactivity, and retriggers if additional movement is detected within 27 seconds. This results in videos of varying lengths, with a maximum length of 30 seconds (however, we deployed two video cameras with a static video length of 30 seconds per trigger). We surveyed 15 camera sites within the range of the Jicarón tool-using group, the majority of which (10) targeted anvil sites. We also surveyed 18 other camera sites in non-tool use areas on Jicarón. We can reliably identify most members of the tool-using group (due to their unique tool-using behavior and increased sampling and coding efforts), allowing us to be confident about which cameras were placed in the tool-using group range. However, other cameras capture multiple non-tool-using groups due to their spread across the island (see Supplemental Tables S1 and S2 for details on camera trap deployments).

58 cameras (39 stills and 19 videos) were deployed during the accumulated sampling period at 33 sites. Out of these deployments, 24 had a single sampling period. The remaining 9 sites were repeatedly sampled (ranging from 2 to 6 deployments in the same location). Average duration of sampling nights per camera was 95.21 (range 9-241), totaling 3 701 sampling nights for the tool-using group and 1 821 nights for non-tool-using groups. Camera traps were deployed at various distances from the coast (0.8 m – 42.9 m for the tool-using group and 0.5 m – 39.2 m for non-tool-using groups). Few camera traps were placed on the beach directly targeting the intertidal zone due to challenges in mounting (i.e., tree availability and seawater) and vandalism.

#### Coding of images & data processing

Still-images were compiled into sequences based on the time between triggers: all bursts of images triggered less than 30 seconds apart were considered part of the same ‘sequence’. Each video of 30 seconds was considered a single sequence. All sequences were coded in Agouti, an online platform for archiving and annotating camera trap data (Casaer et al., 2019). In each sequence, we identified the animal species visible and the number of individuals per species. For analyses, we only considered sequences with capuchins in them (n = 13 728). We did not include any 0’s in our models because an absence of capuchin detections does not necessarily mean an absence of capuchins (e.g., camera traps do not always trigger fast enough to capture a traveling animal). Our analyses thus focus on comparing differences in patterns of detection rates in capuchin activity rather than absence or presence of capuchins. Further, our focus is not to estimate activity patterns as is commonly done for multiple species, using timestamps from camera trap pictures (Ridout & Linkie, 2009; Rowcliffe et al., 2014; Sollmann, 2018). Instead we are modeling the non-random, spatiotemporal patterns of animal detection, which we refer to as *capuchin activity*. We excluded deployment setup and collection days from all analyses, as on these days the human presence may have altered capuchin behavior.

Tidal data, i.e., timing of low and high tides, were obtained from http://www.tide4fishing.com for Cébaco island, which lies ∼90 km from Jicarón, and is used locally by fishermen in the area and digitized into a .csv file using the software Tabula. As the intertidal zone is maximally exposed around the peak of low tide, we calculated the time difference between the initial timestamp of each photo sequence and the time of the nearest low tide. These values ranged from approximately −6 to 6 hours, with 0 indicating the peak of low tide and around −6 and 6 the peaks of high tides. Negative values indicate times where the tide is receding, positive values indicate that the tides are approaching the coast.

We calculated the distance of each camera site to the coast by taking the distance from a camera’s GPS point to the nearest coastal vegetation boundary. To determine the coastal vegetation boundaries, we used high resolution satellite imagery from Planet Labs (Planet Team, 2022). We used 3 m resolution, four-band surface reflectance Planet imagery from January 29, 2021. On this date, cloud cover was near zero and the image was collected at low tide. We used a normalized difference vegetation index (NDVI) threshold of 0.7 to determine the boundary between coastal vegetation and sand or rock. Data were processed in Google Earth Engine and using the terra package in R (Hijmans, 2022). We split the data into dry and wet season based on known rainfall and temperature differences, with the months December-April being part of the dry season and May-November being the wet season.

### Statistical Analyses

We used hierarchical generalized additive models (GAMs) fit using Bayesian regression modeling with Stan via the ‘brm’ function in the brms package v. 2.16.1 (Bürkner, 2017). All statistical analyses were done in R v. 4.2.2 (R Core Team, 2022). GAMs are extensions of generalized linear models that allow estimation of non-linear patterns in data without any prior knowledge of the shape of the expected relationship. Inference is done on the basis of a sum of smooth functions, which are penalized regression splines (Pedersen et al., 2019; Wood, 2006). GAMs are especially useful for seasonal and cyclical data where relationships are unlikely to be linear, such as capuchin activity and tidal cycles. While GAMs are well-suited for identifying non-linear patterns in data, they require caution when used for forecasting based on new data (Pedersen et al., 2019).

We fit five different GAMs: three including capuchin activity in relation to time to nearest low tide and two considering diurnal activity. For the tidal models, we ran an initial model (model MT_1) on all of the data comparing the effect of tidal cycles on activity between tool-using and non-tool-using groups. Then we considered differences in tidal pattern between the dry and wet season for the tool-using (MT_2) and non-tool-using groups (MT_3) separately. The diurnal activity GAMs examined how spatial spread of activity patterns varies depending on the time of day, also separately for the tool-using (MD_1) and non-tool-using groups (MD_2).

#### GAM specifications

To compare tool-users to non-tool-users (model MT_1), we fit a Poisson GAM. Our outcome variable was the count of unique capuchins annotated in a sequence. For predictors, we estimated the effects of i) temporal difference from nearest low tide, ii) distance of camera trap from coast, and iii) an interaction of temporal difference from nearest low tide and camera distance from coast. We used tensor product smooths for this interaction, which allows one to model responses of the outcome variable to interactions of multiple variables with different units.

We used a cyclic cubic spline for time to nearest low tide, as 6 hours before a given low tide matches up to 6 hours after the previous low tide. We also estimated varying smooths of this tensor product for the tool-using and non-tool-using groups, and included if a group were tool-users (1) or not tool-users (0) as an index variable. As we have uneven sampling between the tool-using and non-tool-using group, we include a global tensor smoother for distance to coast and time to low tide, and a tensor smoother considering the tool-using group and non-tool-using groups separately as smooth deviations from that overall surface. We included camera trap location as a random effect.

Secondly, to assess seasonality in a possible tidal effect, we fit two more GAMs with the same structure: one for the tool-using group (MT_2) and one for the non-tool-using groups (MT_3). In these models, we estimated varying smooths of the tensor product for the wet and the dry season, and included the season, wet/dry, as a fixed effect.

Lastly, to investigate the relationship between coastal activity and temperature, we used time of day as a proxy, since temperature varies depending on the time of the day. We used a proxy rather than the actual temperature as camera trap measurements of ambient temperature are unreliable and we lack independent measurements (e.g., from a weather station) to validate them. As such, we fit the same structure GAMs as for the seasonality question, one for the tool-using group (MD_1) and one for the non-tool-using group (MD_2). However, we used hour of the day (from 0-23) rather than time to low tide.

We z-transformed time to low tide, distance from coast, and hour of the day to improve computational speed, model fit, and ease parameter interpretation. All models were fit with mild regularizing priors, using normal(0,2) for the intercept and estimates, and exponential(1) for standard deviations. We performed a prior predictive simulation to visualize the priors. We ran the final models with 3 chains, each having 4000 iterations, including a warm-up period of 2000 iterations per chain. Our model was stable with large effective sample sizes (Bulk_ESS and Tail_ESS over 1000 for nearly all estimates) and Rhat values smaller than 1.01. For all models, Pareto k estimates were below 0.5. We used the posterior predictive check function to visually assess model fit and confirm our choice of priors. For full model specifications and details see the Supplementary Materials and associated reproducible R code.

#### Assessing reliability of model estimates

Tensor-smooth products were visualized into 2D heatmaps (filled contour plots) using the ggplot2 package v. 3.3.6 (Wickham, 2016). To assist in interpretation of these patterns, we used the ‘method of finite differences’ to estimate the first derivative of the spline, which allows for identification of periods of change along a fitted spline. Previous implementations of the method of finite differences on GAMs, were done on single-dimension splines in a frequentist context (Curtis & Simpson, 2014; Monteith et al., 2014). We have built upon this by developing a Bayesian extension of this method for two-dimensional interaction splines (Alavi, 2022). To accomplish this, we first used the ‘posterior_smooths’ function in ‘brms’ to obtain posterior predictions. Then we recomputed posterior predictions after adding or subtracting a small offset, 0.001, for both predictors in the tensor smooth. As predictors were z-transformed, 0.001 represents 1/1000^th^ of one standard deviation for each predictor, a comparable change in both predictors despite their different scales. The first derivative of the two-dimensional tensor smooth can be approximated as follows:

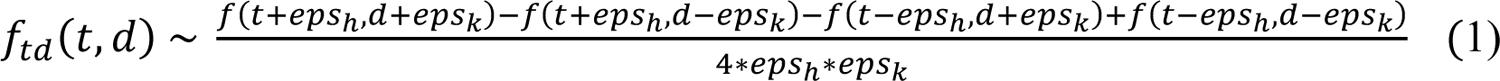

*t* and *d* represent the two predictors in the tensor smooth interaction (e.g. time to low tide and distance to coast). *eps_h_* and *eps_k_* represent the small offset that is added or subtracted to each estimate. With the first derivative approximation, we could identify regions of the 2D surface where the slope was non-flat in all four directions represented in the numerator of equation 1. If 89% of the posterior uncertainty interval of the first derivative was on one side of 0, then we interpreted it as reliable evidence for a non-zero rate of change. This conservative criterion allows us to identify areas of the 2D surface of model estimates where we have the most evidence that: *i)* the model reliably predicts a non-zero change *ii)* this change is consistently (>89% of the time) in the same direction even with small perturbations to the model. Interactive 3D plots of the derivatives of each tensor plot and contourplots showing the proportion of the derivatives above and below zeros are provided as Supplementary Material.

All data and code used for analyses in this paper are available via https://doi.org/10.5281/zenodo.7468017.

## Results

Capuchins live at high densities on Jicarón, but use different locations with varying intensity. We observed capuchins at least once on every camera trap within each deployment (sequence range: 6 - 1224). On average, 2 ± 1.64 (range 1-22) capuchins were observed per sequence. All models indicated considerable variation between camera locations (see Supplementary Figure S4, S6 & S8 for model estimates of capuchin activity per camera location).

### A guide to interpreting 2D heatmaps of derivatives

We present the best predictions of our model in Figure 1. We guide the reader through how to interpret these types of graphs using Figure 1 as an example. In these figures, the y-axis represents the distance from the coast in meters. The x-axis represents hours until and after nearest low tide, where 0 indicates low tide and the boundaries of the graph around −6 and 6 represent high tides. Observations are indicated in the margins of the axes by translucent gray hash marks. In these heatmaps, color reflects capuchin activity: 0 is the mean number of capuchins from z-score transformed raw data. Light colors indicate a greater number of capuchins than the mean, while dark colors indicate a lower number of capuchins. For example, in Figure 1b, the bright yellow peak at 25m and 3.8 hours after low tide indicates highest capuchin activity. The small dark purple trough at 10m from the coast and 5 hours after low tide indicates lowest capuchin activity. We use color saturation to represent evidence for a reliable non-linear change in the number of capuchins as represented in Equation 1. Color saturated areas have more than 89% of the derivative being non-zero, indicating that estimated changes in capuchin activity are consistently in the same direction (i.e., positive or negative).

**Figure 1:**
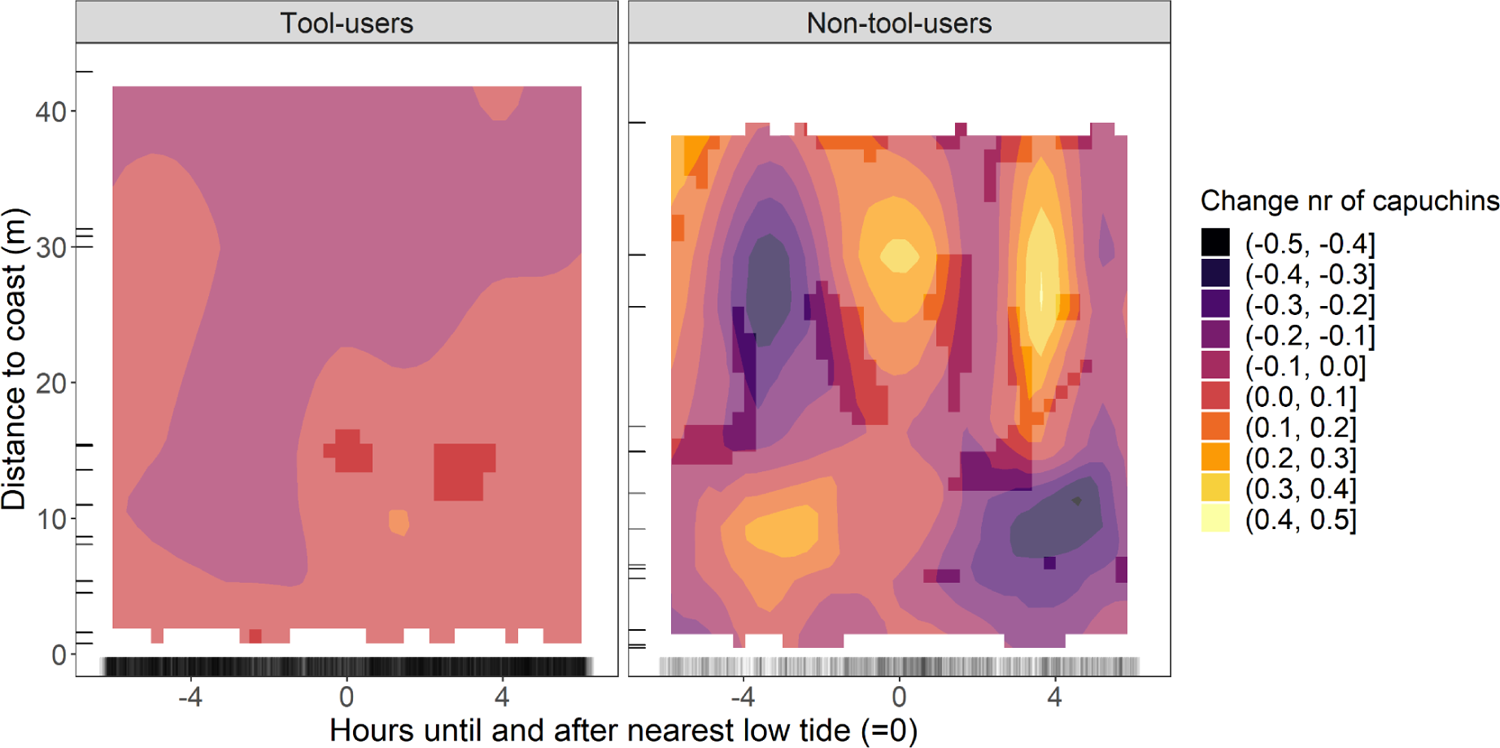
Tidal activity of tool-using vs non-tool-using capuchins. 2D heatmap showing capuchin activity (color) at various distances to the coast (y-axis) and hours until and after nearest low tide (x-axis), for the a) tool-using group and b) non-tool-using groups separately. More color-saturated areas indicate where > 89% of the posterior distribution of the derivative of the tensor smooth interaction lies on one side of 0.

### Tool-users vs. non-tool-users (Model MT_1)

The influence of tides on space use patterns differed for tool-using and non-tool-using groups. While we observe little difference between tool-users and non-tool-users in the number of capuchins present per sequence (tool users: 1.72 [95% CI 1.45-2.08], non-tool-users: 1.75 [95% CI 1.20-2.59]; see Table S3 for estimates), the patterns of capuchin activity relative to the tidal cycle vary.

For the tool-using group (Figure 1a), our model estimates a higher number of capuchins near the coast compared to further inland, although the difference is small (change of <0.2). We see more capuchin detections close to the coast (0-30m) at and after the peak of low tide (0-6 hours) compared to before (−4–0 hours). We see reliable evidence for a change in the number of capuchins in only a small area of the heatmap at 10-15 m from the coast during and 4 hours after the peak of low tide (highlighted in Figure 1a), based on the derivative of the tensor smooth interactions (Figure S5).

In contrast, for the non-tool-using groups (Figure 1b), our model estimates more capuchin activity near the coast prior to the peak of low tide, and less activity after. Estimated inland capuchin activity is highest approximately 4 hours after low tide, and lowest at the same time near the coast. We see evidence for variation in patterns of capuchin activity, with the largest differences occurring at locations further from the coast (>10m).

### Seasonality: Tool-using group (Model MT_2)

We considered seasonality in separate models for the tool-using group and non-tool-using groups. Although for the tool-using group we find comparable numbers of capuchins per sequence between the dry and wet season (dry season: 1.79 [95% CI 1.57-2.08], wet season: 1.82 [95% CI 1.40-2.29]; see Table S4), the patterns of capuchin activity in relation to the tidal cycle differ.

The activity of the tool-using group varies with the tides throughout the year, but stark differences exist between the patterns observed in the dry and wet seasons. In the dry season (Figure 2a), capuchin activity near the coast peaks after high tide (−6 to −2 hours) and is lowest at low tide (0 hours). Changes in when and where capuchins are active are much smaller in the dry season than in the wet season. Based on the derivative of this model there are several combinations of distance and time to low tide where we have reliable evidence for differences in activity (Figure S7). In contrast, in the wet season (Figure 2b), we see the opposite pattern. Our model estimates that capuchin activity is lowest near the coast after the peak of high tide, and is highest near the coast immediately preceding and following low tide (−2 to 2). Additionally, capuchins are most active further inland (20-40m from the coast) immediately after high tide. This is also the time period when capuchin activity at the coast is lowest. The derivative highlights large areas where we have reliable evidence for a change in capuchin activity, concentrated around the peak of low tide and cameras closer to the coast (<30 m).

**Figure 2:**
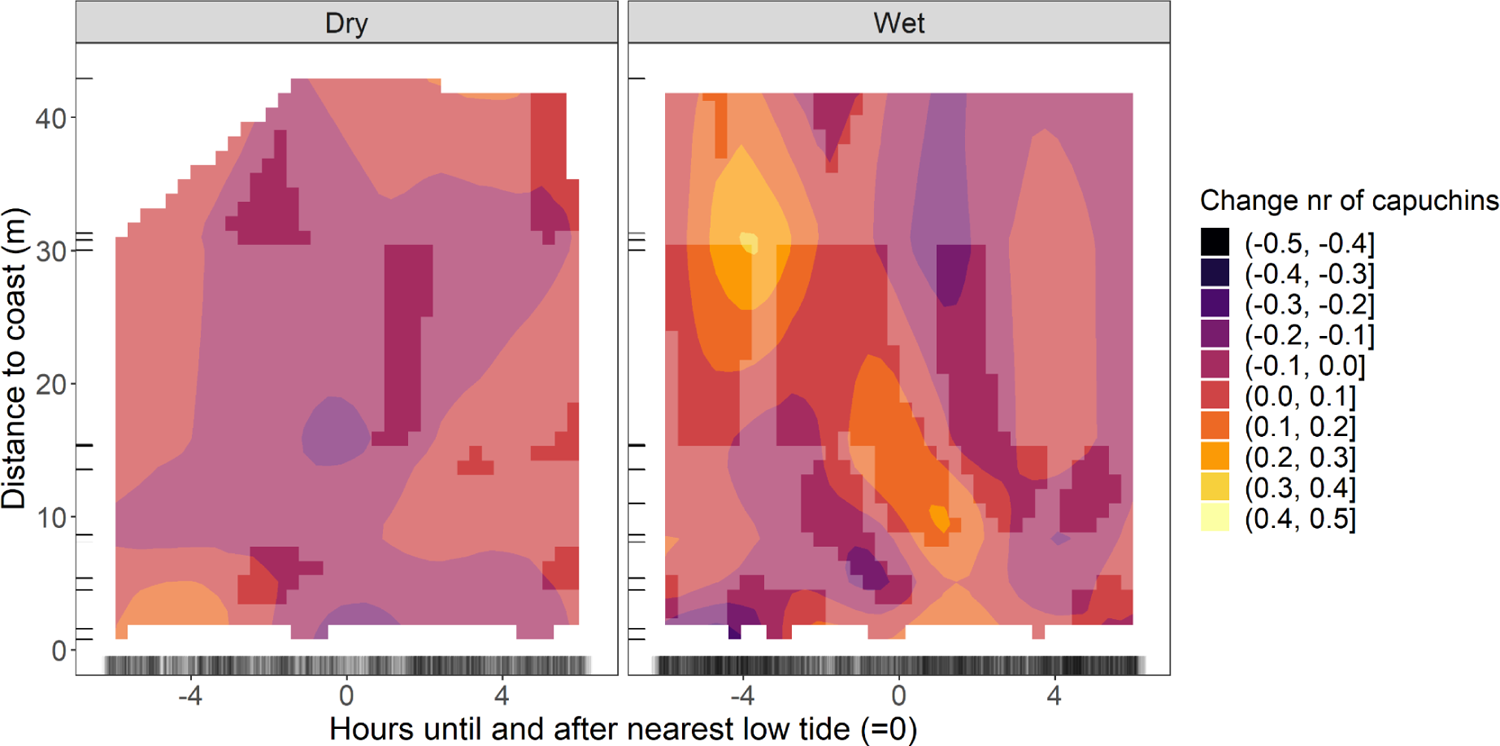
Tidal activity of tool-using capuchins: dry vs. wet season. 2D heatmap showing capuchin activity (color) of the tool-using group at various distances to the coast (y-axis) and hours until and after nearest low tide (x-axis), for the a) dry and b) wet season separately. More color-saturated areas indicate where 89% or more of the posterior distribution of the derivative of the tensor smooth interaction lies on one side of 0.

### Seasonality: Non-tool-using groups (Model MT_3)

Tidal influence on the space use patterns of the non-tool-using groups is most pronounced during the dry season, which is opposite to our findings for the tool-using group, whose activity near the coast is estimated to peak around high tide in the dry season and around low tide in the wet season. As with previous models, we found no change in the average capuchin activity between seasons (dry: 1.79 [95% CI 1.43-2.2], wet: 1.79 [95% CI 1.22-2.56]; see Table S5), but rather changes in the timing of capuchin activity at different distances from the coast.

In the dry season (Figure 3a), the pattern estimated by our model is similar to the predictions of our first model comparing tool-using and non-tool-using groups (Figure 1b). The model estimates higher capuchin activity near the coast in the hours following low tide (0 to 6). However, the peak of capuchin activity as estimated by the model lies after low tide further inland (10-40m). We see reliable evidence for a change in capuchin activity in regions further from the coast (20-40m, Figure S9). In the wet season (Figure 3b), changes in capuchin activity are much smaller than in the dry season. The model estimates a higher number of capuchins near the coast before and during low tide (−6 to 2) than after low tide (2 to 6), however this difference is small (<0.2). There are no regions of the heatmap where we have reliable evidence for a change in activity for non-tool-using groups.

**Figure 3:**
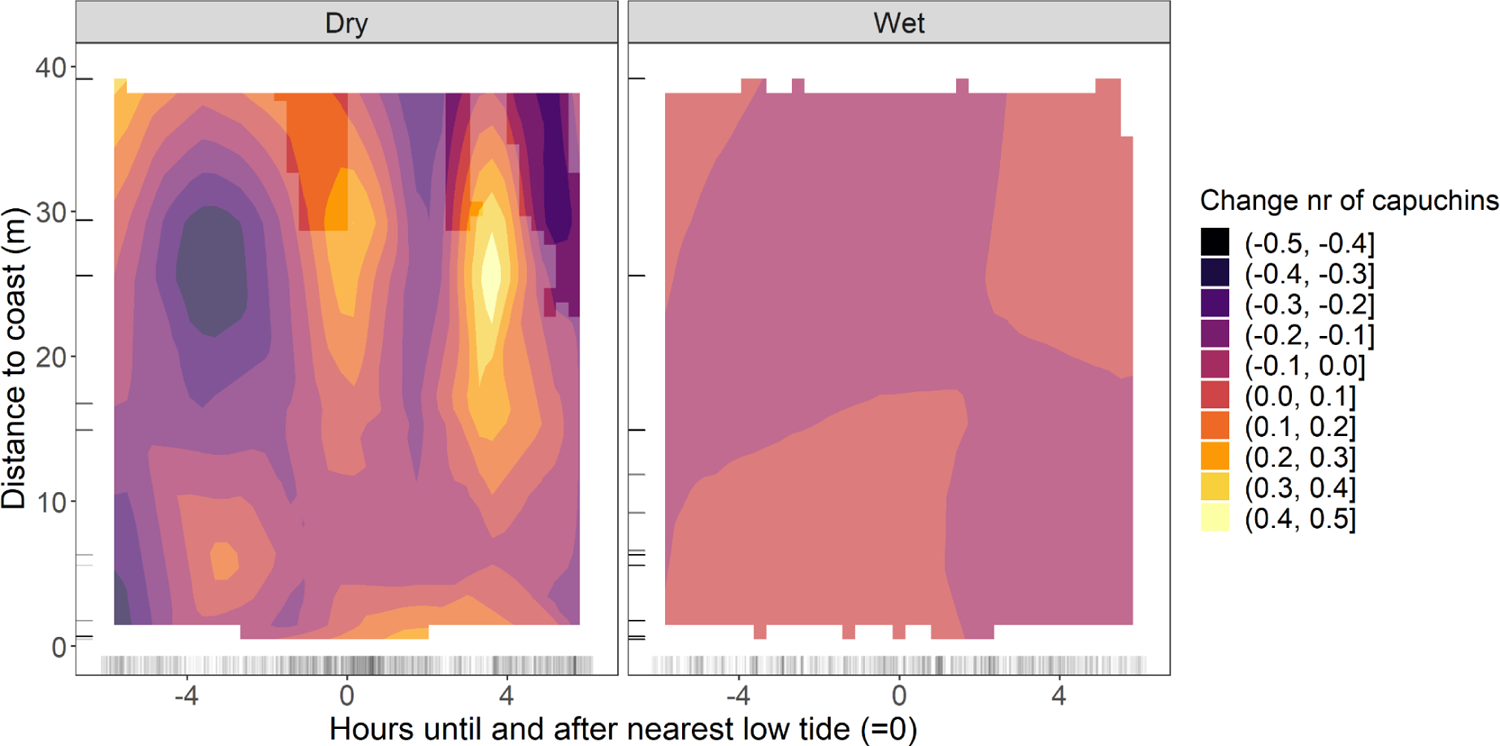
Tidal activity of non-tool-using capuchins: dry vs. wet season. 2D heatmap showing capuchin activity (color) of the non-tool-using groups at various distances to the coast (y-axis) and hours until and after nearest low tide (x-axis), for the a) dry and b) wet season separately. More color-saturated areas indicate where 89% or more of the posterior distribution of the derivative of the tensor smooth interaction lies on one side of 0.

### Daily Activity: Tool-using group (Model MD_1)

Consistent daily shifts in temperature or capuchin movement patterns are possible explanations for varying capuchin detection rates at the coast relative to the tides. For the tool-using group, we see a consistently different activity pattern between the dry and wet season in relation to distance to the coast and time of day.

In the dry season (Figure 4a), our model estimates that capuchin activity near the coast is lowest in the morning, and higher only after 17:00 (see Table S6 for model estimates). Based on the derivative, we have reliable evidence for a change in capuchin activity in several regions, although mostly inland (Figure S10). In the wet season (Figure 4b), we see high coastal activity throughout the day, although the peak of activity lies inland (15-30 m) between 10:00 and 18:00. In the dry season we observe capuchin activity until later in the day (21:00) than in the wet season (19:00). In several regions of the heatmap we have evidence supporting a change in the number of capuchins.

**Figure 4:**
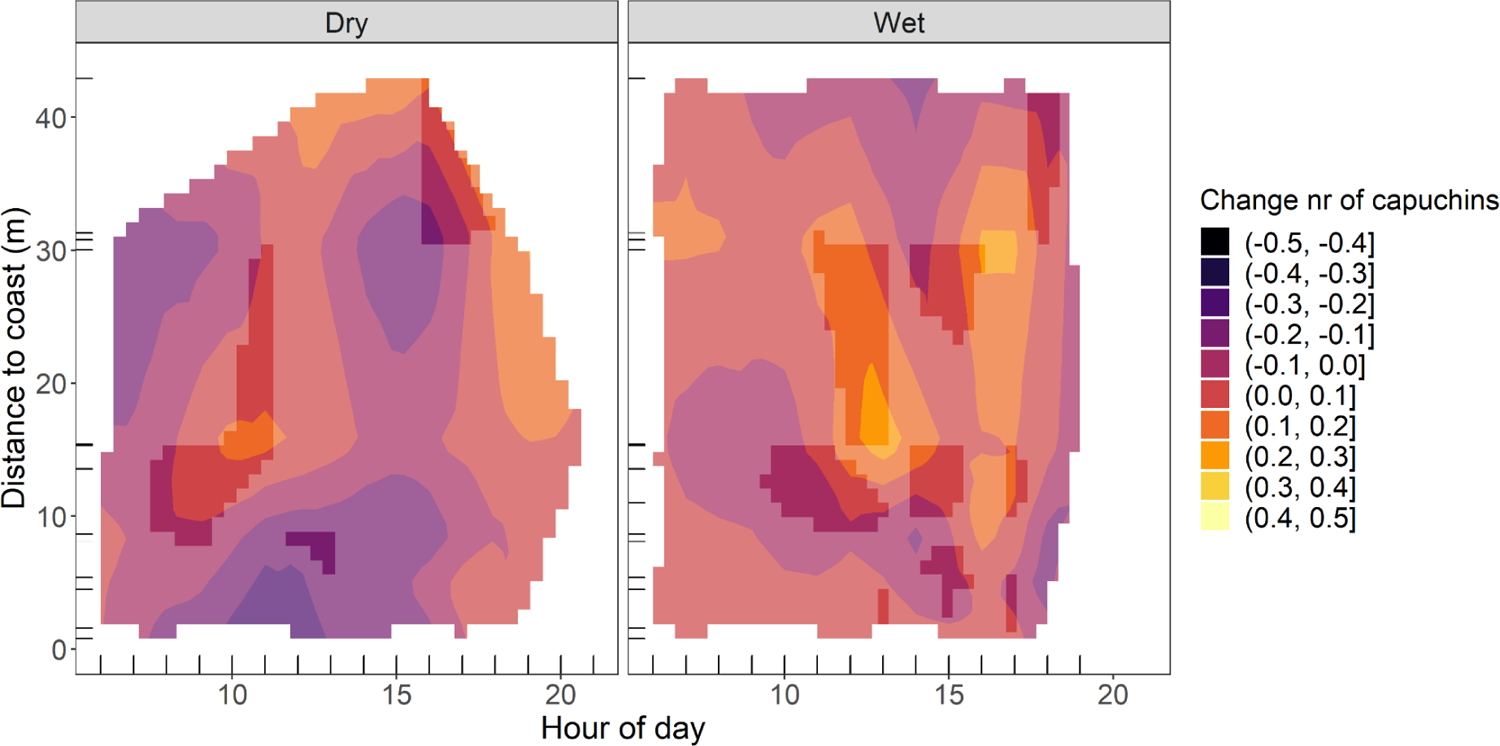
Diurnal activity of tool-using capuchins: dry vs. wet season. 2D heatmap showing capuchin activity (color) of the tool-using group at various distances to the coast (y-axis) and hours of the day (x-axis), for the a) dry and b) wet season separately. More color-saturated areas indicate where 89% or more of the posterior distribution of the derivative of the tensor smooth interaction lies on one side of 0.

### Daily Activity: Non-tool-using groups (Model MD_2)

For the non-tool-using groups, we observed an opposite pattern to the tool-using group when considering activity at varying distances to coast in relation to time of day (Figure 5, see Table S7). In the dry season (Figure 5a), our model estimates highest capuchin activity inland (30-40 m) in the morning until 13:00. Estimated capuchin activity is lowest in the afternoon (13:00-18:00) from near the coast until 30 meters inland. The model estimates a higher coastal activity in the morning, but only for a small area of the 2D heatmap 30-40m inland at 16:00-17:00 do we have reliable evidence for a change (Figure S11). In the wet season (Figure 5b), estimated changes in capuchin numbers are considerably smaller than in the dry season, and there are no areas of the 2D heatmap where the derivative provides reliable evidence for a change. The model estimates a slightly higher number of capuchins near the coast than further inland throughout the day, with highest activity in the late afternoon from ∼16:30 onwards.

**Figure 5:**
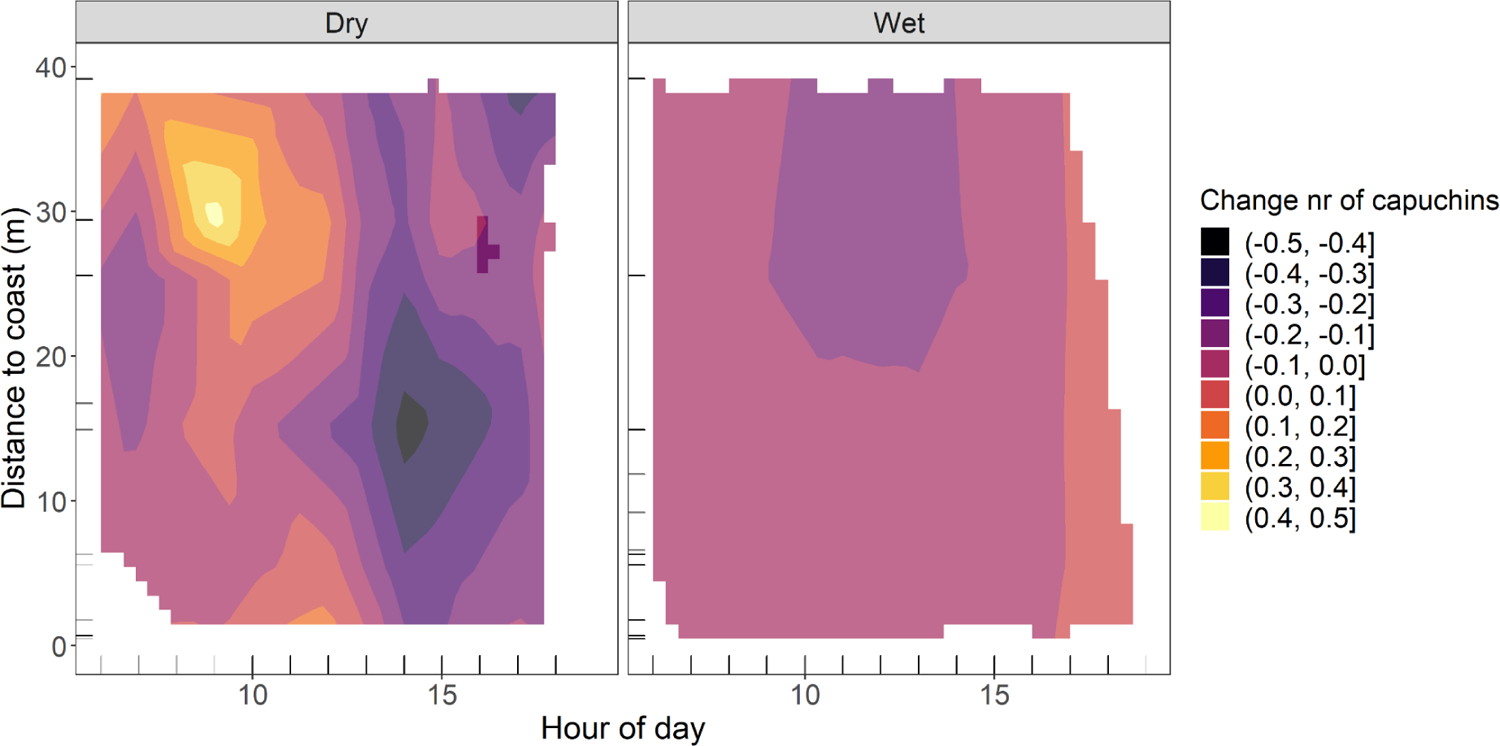
Diurnal activity of non-tool-using capuchins: dry vs. wet season. 2D heatmap showing capuchin activity (color) of the non-tool-using groups at various distances to the coast (y-axis) and hours of the day (x-axis), for the a) dry and b) wet season separately. More color-saturated areas indicate where 89% or more of the posterior distribution of the derivative of the tensor smooth interaction lies on one side of 0.

## Discussion

White-faced capuchins living on Jicarón island adjust their activity patterns in correlation with tidal cycles. Our findings are consistent with the hypothesis that availability of intertidal resources plays an important role in shaping patterns of movement and habitat use. We find seasonal variation in the relationship between capuchin activity and the tidal cycle, as well as in capuchins’ daily activity patterns. Tool-using and non-tool-using capuchins are seen less frequently near the coast in the dry season during the hottest times of the day (∼13:00-17:00), suggesting that high temperatures likely limit capuchins’ activity near the coast and potentially in the intertidal zone. Although capuchins in both tool-using and non-tool-using groups consume intertidal resources, they show different patterns of activity in relation to tidal cycles. Activity of the tool-using group near the coast increases around high tide in the dry season, and around low tide in the wet season. Patterns are more pronounced in the wet than the dry season, when differences in estimated capuchin activity are larger and we have more reliable evidence supporting these changes. In contrast, non-tool-using groups’ activity near the coast is higher around low tide in the dry season and around high tide in the wet season, with more pronounced differences in capuchin activity in the dry season.

### Capuchin activity varies in relation to tidal cycles

By showing that tool-using and non-tool-using capuchins’ activity at the coast is related to the tidal cycle, we provide additional evidence that exploitation of intertidal resources by maritime mammals shapes the rest of their activity patterns, similar to seabirds (Irons, 1998) and arctic foxes (Nielsen, 1991). While from our data we cannot conclude whether exploitation of the intertidal zone is opportunistic or obligate (Carlton & Hodder, 2003), the tight coupling between Coiban capuchin’s coastal activity and tidal cycles indicate that Coiban capuchins likely exploit intertidal resources regularly. Tool-using and non-tool-using capuchins consistently spend time near the coast at either low or high tide, depending on the season. Because the timing of low tide shifts each day, these patterns cannot arise from intermittent activity near the coast irrespective of the tidal cycle. These findings indicate that Coiban capuchins are timing their activity at the coast to coincide with specific parts of the tidal cycle.

### Temperature limits coastal activity

Tool-using and non-tool-using capuchins show increased coastal activity at different periods of the tidal cycle (Figure 1). To understand these differences, we first have to consider differences in coastal activity between seasons. If seasons are jointly analyzed, signatures of seasonal variation in coastal activity can be weakened or masked if a pattern is only present in one season and absent in the other, or if seasonal patterns are opposite to one another. Higher temperatures and lack of rainfall in the dry season affect capuchins’ daily activity and, potentially, their exploitation of the intertidal zone. During the hottest times of the day, capuchins prefer resting rather than energy-expending activities like foraging and traveling (Campos & Fedigan, 2009), and presumably, tool-use. This is supported by both the tool-using group and non-tool-using groups showing lower activity in the dry season during the hottest time of the day (∼15:00) at all distances to the coast (Figures 4a & 5a). Capuchins might compensate for missed daylight activity by staying active later in the evening. We see this for the tool-using group who shows activity up to two hours later in the dry season than in the wet season (Figure 4b).

Temperature regulation might also be a driving force to spend time near the coast in general, or at varying moments of the tidal cycle. Overall, the microclimate at the coast is cooler than inland, due to the atmospheric phenomenon of sea breezes which carries cool air to the coast (Kuwagata et al., 1994). At high tide, this cooler microclimate would be more pronounced than at low tide, when the water has further receded. While hot temperatures might push capuchins to the coast where it is cooler, heat might also repel them from the intertidal zone. When it is hot, capuchins might be less inclined to venture into the intertidal zone, where there is no shade, the rocks and sands heat up in the sun, and water in tidal pools and crevices evaporates more quickly. This rapid evaporation increases water temperature and salinity, which reduces the number of species that survive in tidal pools. Thus, at high temperatures, capuchins might be attracted to the cooler coast, but avoid the exposed intertidal zone where it is hotter with less available prey.

### Seasonality in capuchin activity: tool-users vs. non-tool-users

The tool-using group is only consistently close to the coast in the dry season right after high tide (Figure 2a), which supports the idea of temperature limiting their (coastal) activity. Immediately after high tide, the sand is still wet and cool from the receding water and the capuchins can remain close to the shade of the tree line, making it possible to consume snails, such as *Nerita scabricosta,* which migrate to the top of the intertidal zone at high tide (Levings & Garrity, 1983). In the wet season, when temperature is not a limitation, we see high coastal activity of the tool-using group throughout the day (Figure 2b). This group may exploit the full intertidal range that is exposed in the hours before and after low tide. In contrast, the non-tool-using groups show highest coastal activity in the dry season around low tide (Figure 3a), which is unexpected if temperature is a major limitation.

However, when considering their activity at low tide at all distances from the coast we observe more capuchin activity inland than at the coast. One possible explanation is that non-tool-using capuchins are at the coast during low tide only for some low tides of the day, while for others they are resting or further inland. Some low tides will be cooler than others: the intertidal zone is colder during a low tide at 9:00 than one at 14:00. Non-tool-using capuchins might be near the coast in the dry season when it is not the hottest time of the day, and only venture into the intertidal during these cooler low tides, which would lead to the observed pattern. The non-tool-using groups do not appear to consistently be closer to the coast around low tide, but rather only after high tide (Figure 3a). Additionally, differences in capuchin activity are rather small and less certain than for the tool-using capuchins. This indicates that non-tool-using capuchins show less consistent activity patterns with the tidal cycles.

### Seasonality in food availability as a driver of intertidal exploitation

Despite temperature limiting activity in the dry season, this appears to be when non-tool-using capuchins consume most intertidal resources as they show the strongest tidal pattern (Figure 3a). The contrasting pattern of tool-using capuchins showing most coastal activity correlated with the tidal cycle in the wet season raises the question: what role does seasonality in food availability play in intertidal exploitation? Resource scarcity is likely experienced differently by tool-using and non-tool-using groups, as tool-using groups can access structurally protected resources more efficiently (Biro et al., 2013). We have no direct information on seasonal fluctuation in food availability on Jicarón, but tool use occurs more frequently in the transition periods between wet and dry seasons, which might be in response to a limitation in terrestrial resources (Barrett et al., 2018). It is important to consider that capuchins are dietary generalists who not only consume fruits but also insects and other invertebrates, which have their own seasonal fluctuations. Additionally, oceans are also seasonal and, as such, resources in the intertidal zone likely vary spatially and temporally across seasons (e.g., snails, Collin et al., 2017). However, for tool-using capuchins, exploitation of intertidal resources might not be driven by a scarcity of terrestrial resources: in the wet season there is likely peak availability of one of the tool-users most consumed resources, *Terminalia catappa,* which bears fruit from January to April and from May to September (Perez & Condit, n.d.). This coincides with high rates of coastal activity around low tide and the possible exploitation of intertidal resources. Further, tool-using capuchins not only consume fresh sea almonds, but old ones which accumulate in the leaf litter and wash up to shore as well (pers. obs.). Thus, they may be less dependent on the seasonality of tree fruit productivity.

### Tool users vs non-tool-users: why such different patterns?

Why do we observe a difference in the relationship between activity and tidal cycles for the tool-using group and non-tool-using groups? The non-tool-using groups show more activity at the coast during low tide in the dry season, when temperature might also play a big role, and no clear tidal pattern in the wet season (Figure 3). The tool-using group’s activity is more consistent with exploitation of tidal resources in both seasons though at different moments depending on temperature (high tide in the hot dry season and low tide in the cooler wet season, Figure 2). We propose two explanations for why non-tool-using groups might exploit intertidal resources more opportunistically or more rarely than tool-using capuchins.

First, the spatial fixedness of both materials and resources for tool use might drive tool-users to spend more time near the coast. Stones large enough to serve as anvils and hammerstones are more abundant at the coast and the mouth of streams than further inland and may also be prevalent in the intertidal zone itself. Further, important resources, including bivalves, crabs, and sea almond trees are restricted to the coast (Perez & Condit, n.d.). Because of these constraints, the tool-using group’s important foraging spots and anvils are all close to the coast, giving tool-users more opportunities to pick up visual and auditory cues of low tide and more opportunities to exploit tidal resources. Non-tool-using groups, who also forage on sea almonds but only eat the less calorically dense exocarp of ripe fruits, but not the nut inside (Barrett et al., 2018 also see Campbell, 2013 for evidence in another population), likely need to forage more widely for resources and do not have a similar reliance on coastal locations as the tool-using group. Thus, increased proximity to the coast due to reliance on coastal tool use resources might allow and encourage the tool-using group to forage in the intertidal zone more frequently.

A second explanation is that tool use allows the tool-using group to exploit intertidal resources more consistently than the opportunistic foraging by non-tool-using groups. By allowing access to encapsulated resources, percussive tool use can both broaden the diversity of consumable food items, improving diet quality (Izar et al., 2022), and increase the efficiency of consuming known resources (Biro et al., 2013). Both (or either) of these aspects of tool use could explain why the tool-users might show more intertidal foraging than non-tool-using groups. The costs due to energy investment, time lost foraging elsewhere, and risk of venturing out into the exposed intertidal zone might be outweighed by tool-users’ ability to open more resources with greater efficiency in the intertidal than non-tool users.

Differences in sampling between the tool-using group and non-tool-using groups in spatial and temporal coverage (a small area with dense camera trapping vs a larger area with sparse camera trapping) as well as comparing one group to a multitude of groups may have affected our results. This possible limitation implies that while we can be more confident of the presence of a tidal pattern in the tool-using group, we should interpret the tidal pattern in the non-tool-using groups’ results with more caution. We also have less reliable evidence for a change in the capuchin activity for the non-tool-using groups’ models than for the tool-using groups’ models. Given this limitation, we can tentatively conclude that for coastal-living capuchins, investing in consistent exploitation of intertidal resources might be a trade-off between nutrition gained and energy and time expended in which using tools could be a decisive factor.

### Opportunities for future research

To disentangle capuchins visiting the coast for temperature regulation versus foraging on intertidal resources, we need data on capuchin activity from the intertidal zone itself. With camera traps, this was not possible, thus we infer visits to the intertidal based on proximity to the coast. We know that both tool-using and non-tool-using capuchins forage on intertidal resources from in-person observations (Barrett et al., 2018; BB, MC, CM, ZG pers. obs.), and physical evidence of tool use in the intertidal zone (Figure 6). Additionally, capuchins on the island of Coiba also consume intertidal resources like snails (Méndez-Carvajal & Valdés-Díaz, 2017). However, more than one process could contribute to the patterns of activity we report, for example, capuchins being near the coast to regulate their temperature. We consider this alternative by including spatial patterns depending on time of day, where time of day serves as a proxy for temperature. Our findings that temperature likely constrains capuchin activity near the coast suggest that in future research this should be examined in more detail. One possibility is to directly measure the ambient temperature at varying distances from the coast. Additionally, with semi-arboreal species such as capuchins, terrestrial camera traps miss activity: we cannot distinguish an absence of capuchins from capuchins present but in the trees. While the capuchins on Jicarón are highly terrestrial (Monteza-Moreno, Crofoot, et al., 2020), we attempted to further mitigate this concern by focusing solely on fluctuations in the number of capuchins present in a sequence, rather than drawing conclusions based on absence. We did not place our camera traps randomly, but in targeted locations, such as on anvils, as this study is part of research on tool use. By only comparing fluctuations in capuchin activity *within* a specific camera location we account for this (and other) variation present between different camera locations.

**Figure 6:**
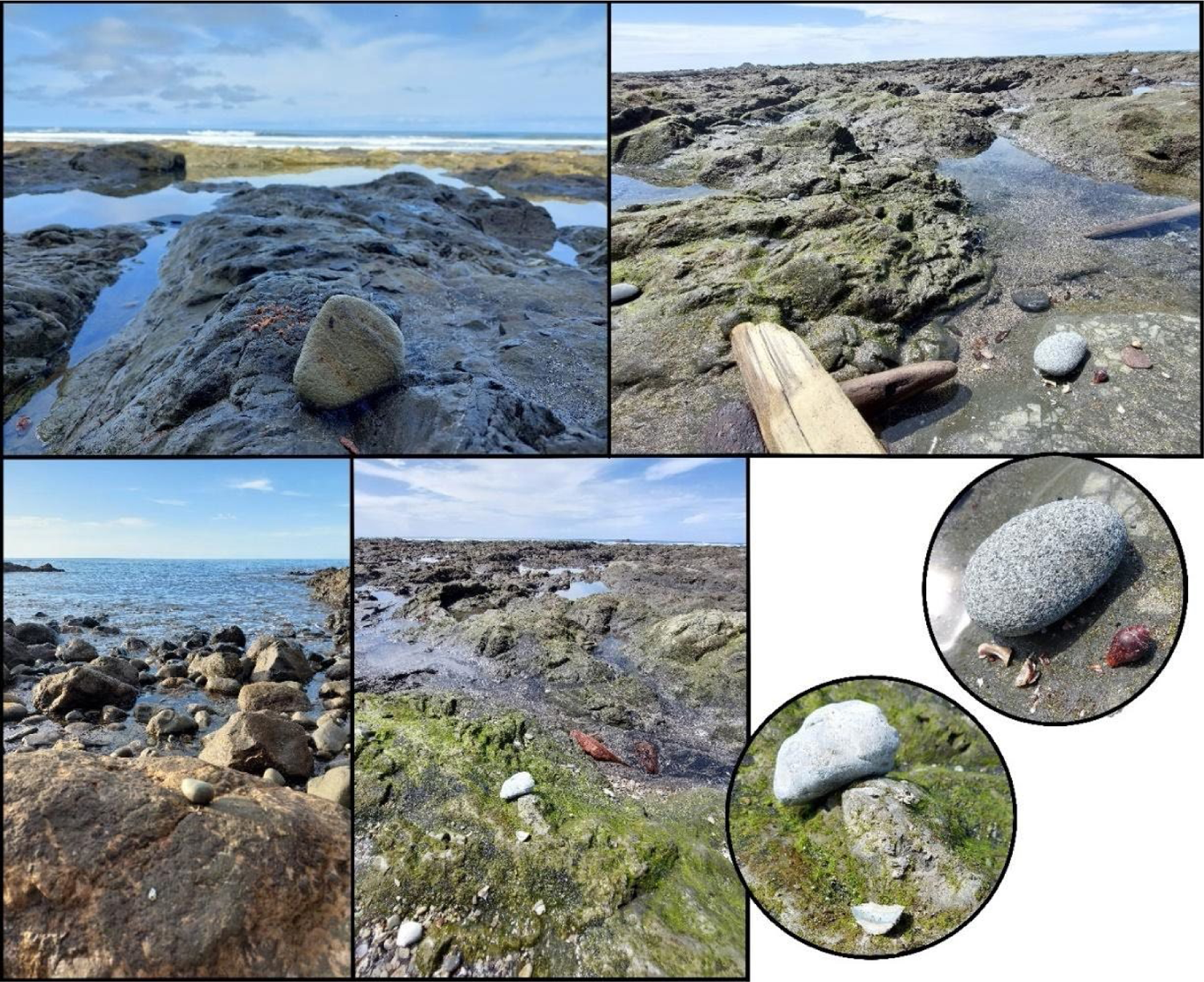
Evidence of tool use in the intertidal zone. Hammerstones and debris in the intertidal zone at low tide, which will be washed away by high tide. Photos by Meredith Carlson & Zoë Goldsborough.

Our findings of such clear tidal patterns in capuchin activity raise the question of how capuchins are aware of the timing of the tides: do they rely on sensory cues (e.g., auditory, visual, or olfactory), cues from other species (i.e., seabird activity), or perhaps even cognitively “track” the tidal cycle? Movement data would provide insights on the exact timing of capuchins’ visits to the coast, and what underlying mechanism(s) allow(s) them to be there at the right time. Additionally, as the spatial scale of this study (0-40 meters from the coast) is limited in comparison to typical capuchin home range sizes (e.g., 0.8-1.5 km^2^ on Barro Colorado Island; Crofoot, 2007), tracking of individuals would also allow for integration of our findings into a more comprehensive understanding of Coiban capuchin space use.

There are additional avenues to examine differences in consumption of intertidal resources by the tool-using group and non-tool-using groups such as DNA barcoding or stable isotope analysis to indicate what proportion of the capuchins’ diet is marine resources (*sensu* Lewis & Sealy, 2018). Additionally, a more detailed examination of the variation in daily activity between the tool-using group and non-tool-using groups is needed to explore the effect tool use might have on their behavior aside from allowing access to tidal resources. Lastly, taking a similar approach in other tool-using and non-tool-using maritime mammals would shed light on whether tool use is indeed important for consistent exploitation of intertidal resources.

### Broader implications of our findings

We found that capuchins living in coastal ecosystems show changes in their activity near the coast that correspond to tidal cycles. While both tool-using and non-tool-using capuchins show tidal patterns that correspond with exploitation of intertidal resources, tool-using capuchins show a stronger pattern in both the wet and dry seasons. Our findings suggest that tidal resources may be very important for coastal-living capuchins, but that tool use might be a prerequisite for their efficient exploitation. Although we lack mainland data for comparison, it is important to consider that Coiban capuchins in this study live in a coastal, *insular* habitat. Islands are often resource-limited: species richness decreases with island area and distance to mainland or insular source populations. Additionally, habitual reliance on tool use is observed in many endemic island-living species (i.e., Woodpecker finches [*Cactospiza pallida;* Eibl-Eibesfeldt, 1961], New-Caledonian crows [*Corvus moneduloides*; Rutz & St Clair, 2012], Hawaiian Crows [*C. hawaiiensis*; Rutz et al., 2016], and Keas [*Nestor notabilis*; Goodman et al., 2018]), or restricted to populations that live on islands (i.e. Burmese long-tailed macaques on islands in the Andaman Sea, Gumert et al., 2009). Potentially, the presence of such a rich intertidal zone in an otherwise challenging ecosystem might be one explanation for why tool use is more likely to arise on islands: to aid exploitation of the intertidal zone.

Percussive tool use on coastal resources also played an important role in human evolution: prehistoric *Homo sapiens sapiens* used stone tools to access a variety of marine resources from the intertidal zone (Erlandson, 2001; Marean et al., 2007; Mazzia & Flegenheimer, 2015). Consumption of marine resources is argued to have aided the rapid increase in brain size observed in human evolution, due to their high contents of docosahexaenoic acid (DHA) as well as iodine and selenium (Crawford & Broadhurst, 2012; Joordens et al., 2014). Modern coastal-living humans do still employ tools to forage in the intertidal zone, and have been found to prefer foods that provide the best nutritious value for the least effort (Mannino & Thomas, 2002). Ethnoarchaeological work in humans has shown that intertidal foraging is a high-reward type of foraging – *if* the tidal conditions are optimal (De Vynck et al., 2016). Nonhuman primates like white-faced capuchins provide an interesting parallel to understand the role tool use has played in human evolution, in the absence of clear evidence from the fossil record.

The findings of this study have implications for the conservation of these capuchins, namely highlighting the potential importance for exploitation of tidal resources for both tool-using and non-tool-using capuchins. Exploitation of intertidal resources by tool-using capuchins appears to be less frequent in the dry season, and due to anthropogenic climate change these periods of hot, dry weather are globally becoming longer and more intense (IPCC, 2022). Further, the ocean is becoming increasingly acidified, which negatively affects many marine organisms (Gaylord et al., 2015) and as such will eventually affect the capuchins too as they forage on these marine resources. The role intertidal zones play in capuchins’ foraging patterns should be included when considering how to best conserve the capuchins on Jicarón and their unique tool use behavior (Brakes et al., 2021).

Overall, our findings both uncover the impact exploitation of tidal resources may have on an animal’s general activity pattern, as well as increase our understanding of how tool use may provide access to a new niche. Animals that are dietary generalists can opportunistically complement their diet with valuable resources from the intertidal zone, and tool use might be necessary to make the investment in these fluctuating resources worth the costs of keeping up with the tidal cycle.

## Ethics Statement

Data for this study was collected as minimally invasively as possible. We obtained ethical permission for this study from the relevant authorities, namely STRI and the Ministerio de Ambiente, Panama (scientific permit no. SC/A-23-17, SE/A-98-19, and corresponding renewals and addenda).

## Acknowledgments

We want to thank the staff at STRI, particularly the field station on Rancheria, for their support and assistance, and Lucia Torrez, Katrin Dieter and Angie Ruiz for supporting the logistics of this research. Additionally, Eliezer Vega, Chris Dillis, Pedro Luis Castillo, Zarluis Miguel Mijango and Evelyn Del Rosario assisted with fieldwork. We are grateful to Lucy Aplin, Edwin van Leeuwen, and Meredith Carlson for their advice on this study, and to Alison Asbury for writing suggestions and feedback.

## Supplemental Material

**Figure S1.**
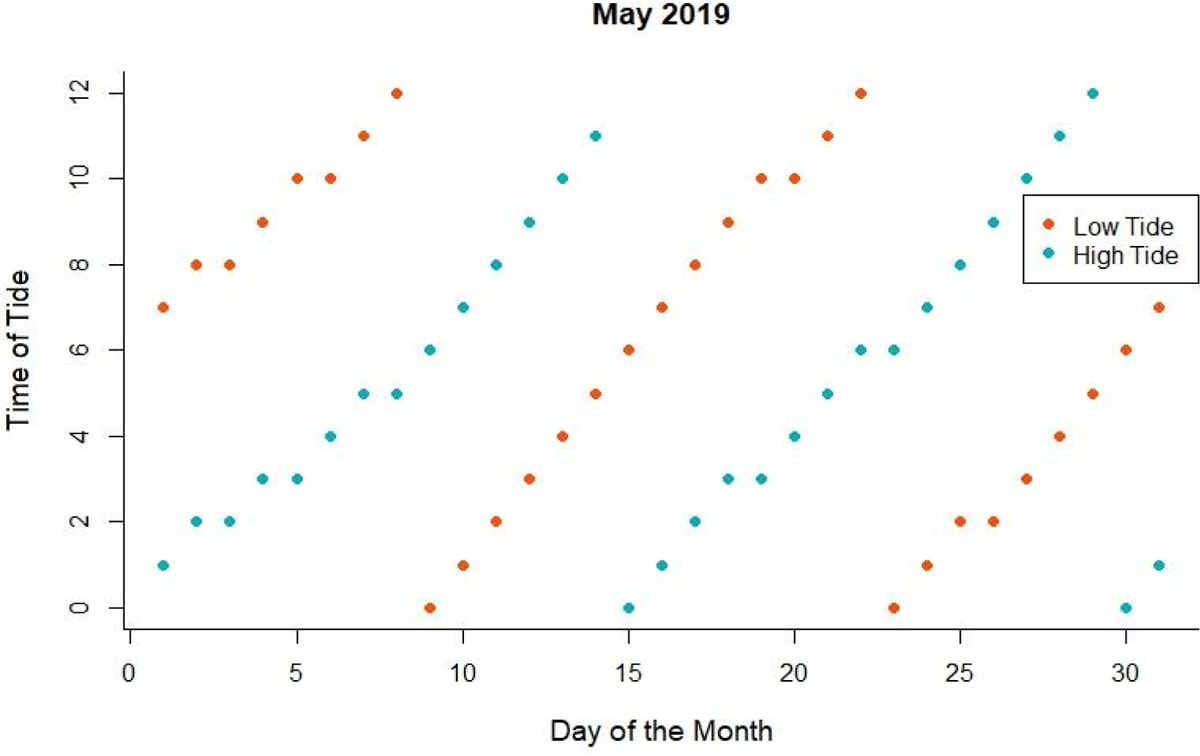
Timing of tides at Jicarón in one month (May 2019)

**Table S1.**
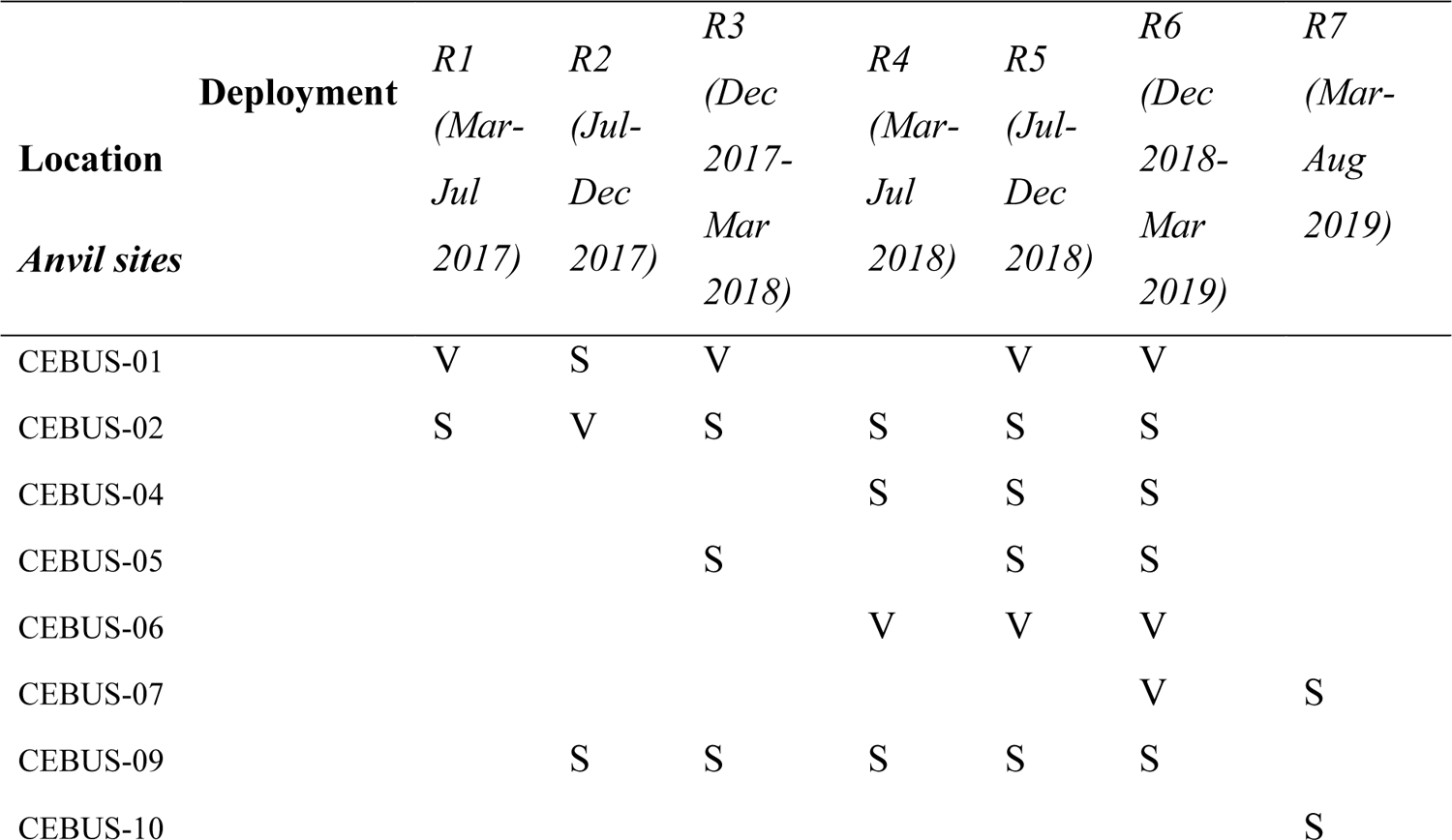
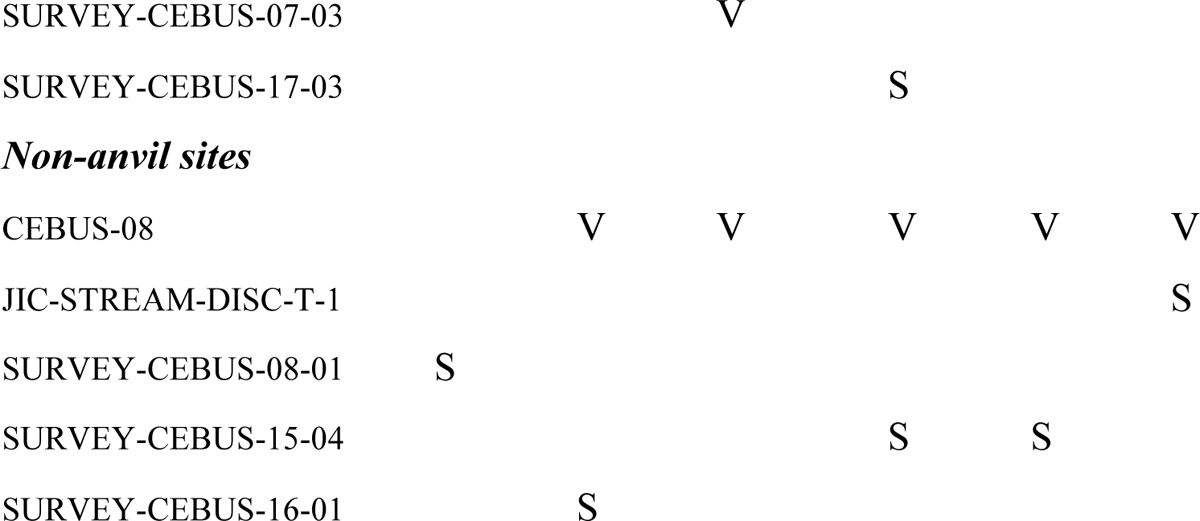
Overview of still (S) and video (V) camera’s deployed for tool-using group

**Table S2.**
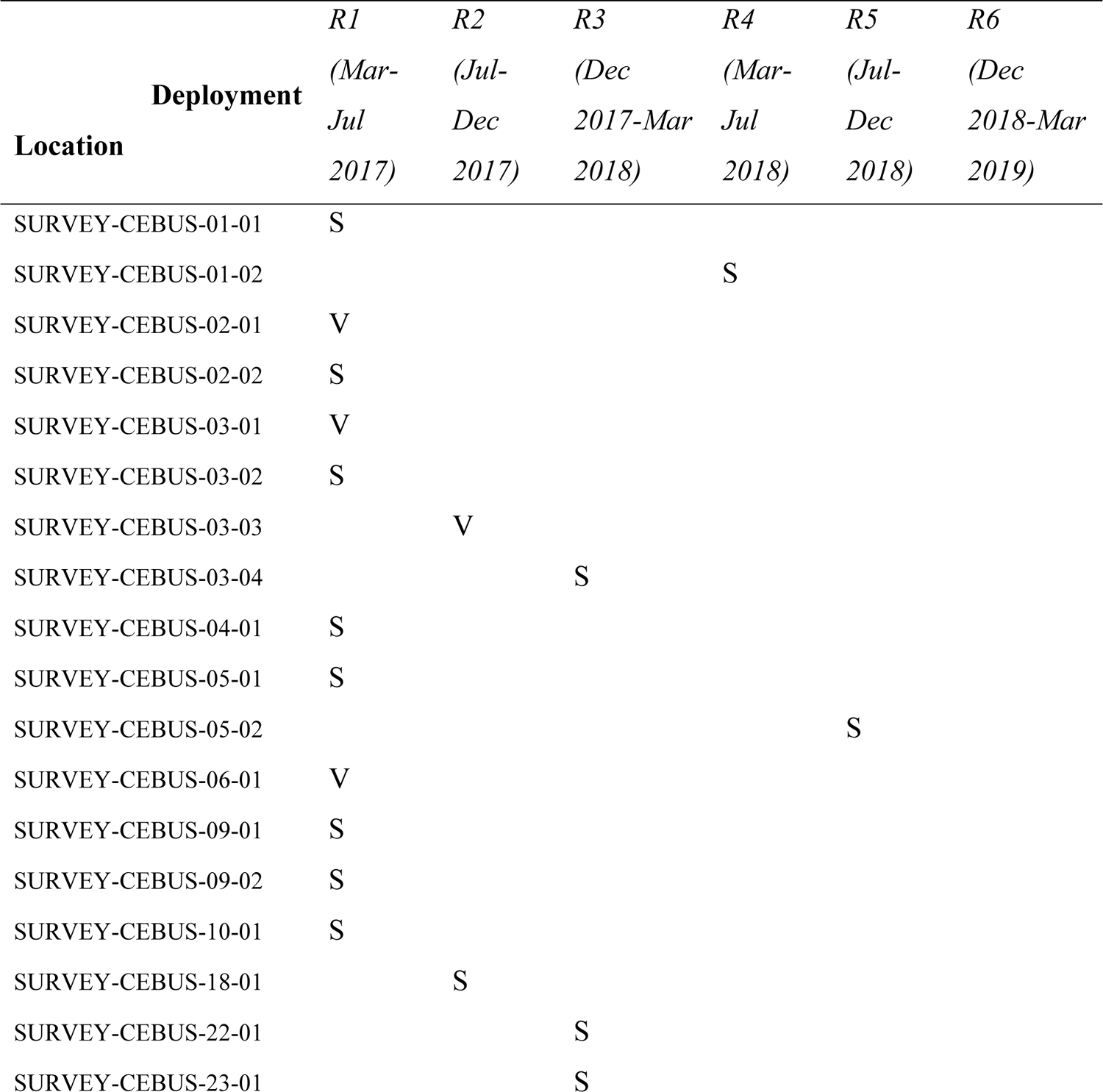
Overview of still (S) and video (V) camera’s deployed for non-tool-using groups

### Model MT_1: Comparing tidal patterns of tool-using group and non-tool-using groups

**Table S3.** Posterior mean model estimates of Model MT_1, a Poisson GAM (Bayes R^2^ = 0.09). All estimated population-level effects are on the logit scale. Non-tool-using groups are the reference category (the intercept).

|  | *Estimate* | *CI_95_low* | *CI_95_high* |
| --- | --- | --- | --- |
| *Smooth terms* |  |  |  |
| Tidetime-Distance_1 | 0.50 | 0.02 | 1.61 |
| Tidetime-Distance_2 | 0.96 | 0.03 | 3.62 |
| Tidetime-Distance_3 | 0.64 | 0.02 | 2.15 |
| Tidetime-Distance_4 | 0.58 | 0.01 | 2.09 |
| Tidetime-Distance non-tool-users_1 | 3.50 | 2.20 | 5.05 |
| Tidetime-Distance non-tool-users_2 | 1.32 | 0.04 | 3.99 |
| Tidetime-Distance non-tool-users_3 | 1.05 | 0.03 | 3.72 |
| Tidetime-Distance tool-users_1 | 0.55 | 0.05 | 1.42 |
| Tidetime-Distance tool-users_2 | 0.94 | 0.03 | 3.08 |
| Tidetime-Distance tool-users_3 | 0.43 | 0.01 | 1.36 |
| Camera location | 0.21 | 0.14 | 0.29 |
| *Population-Level Effects* |  |  |  |
| Intercept | 0.54 | 0.37 | 0.73 |
| Tool-users: Yes | 0.02 | -0.19 | 0.22 |
| Tensor product of Tidetime-Distance | 0.03 | -0.09 | 0.14 |

**Figure S4.**
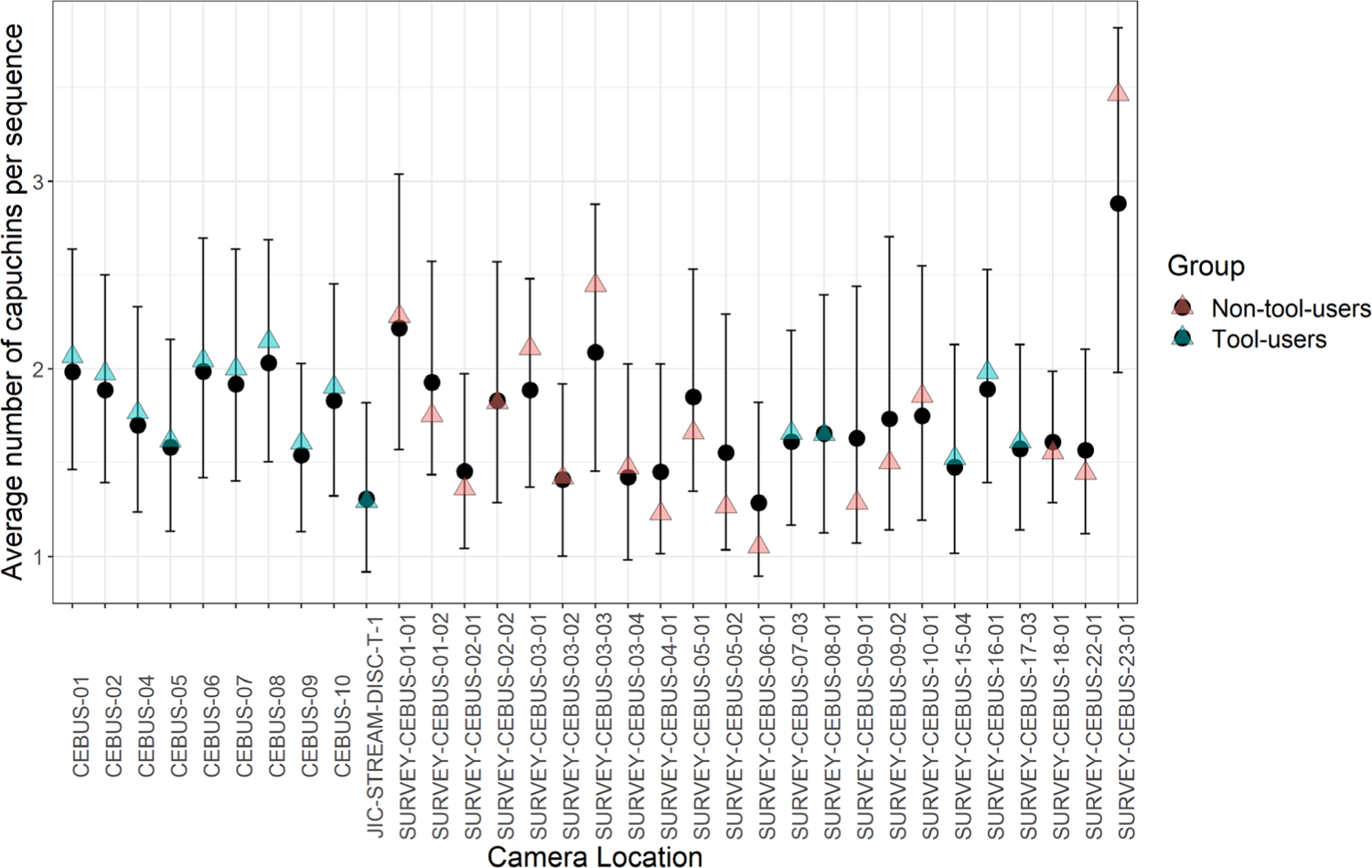
Model MT_1’s posterior mean estimates of the average number of capuchins per camera location represented by black circles. Bars indicate the 95% credible interval. Triangles represent the real average. Blue indicates tool-using and red non-tool-using groups.

**Figure S5.**
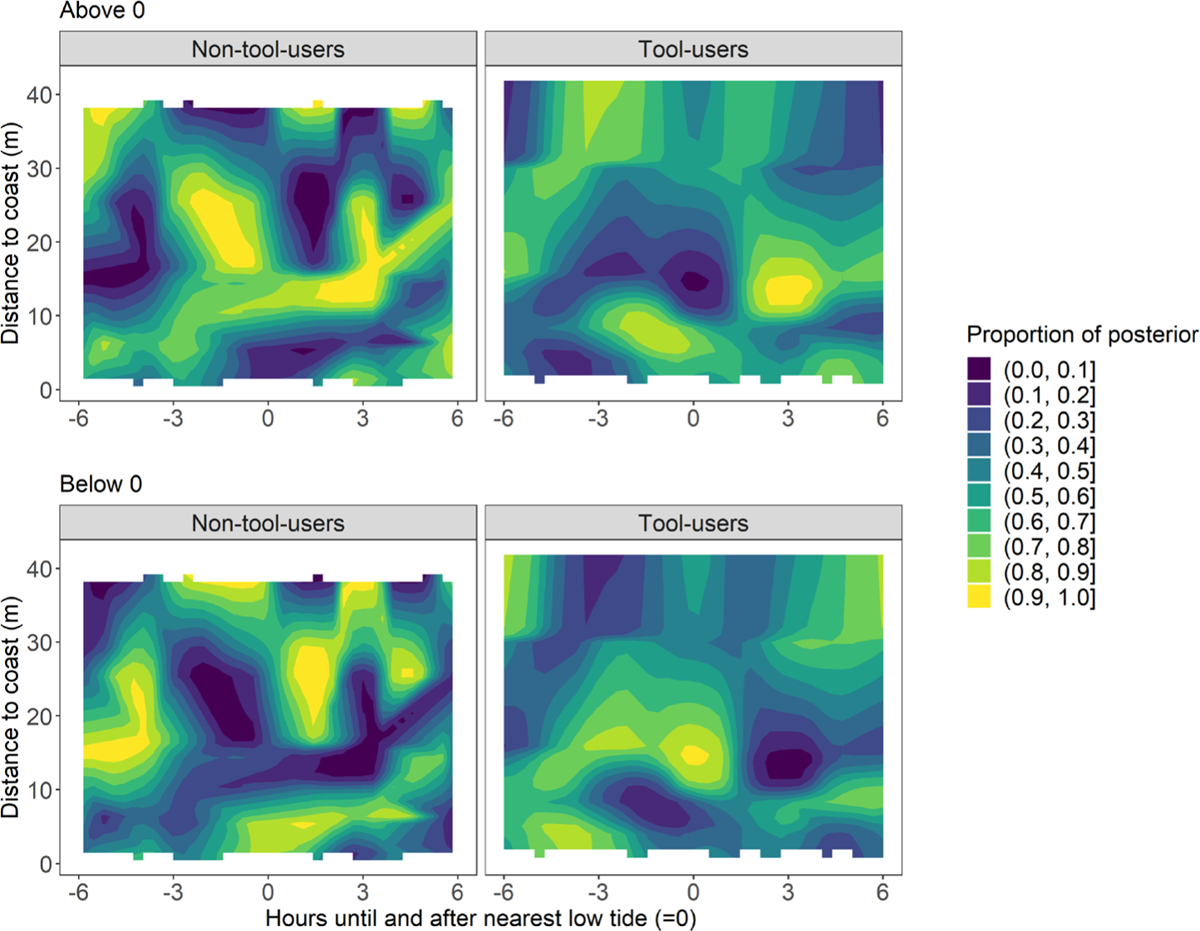
2D heatmap showing for Model MT_1 what proportion of the posterior of the derivative lies above or below 0 as calculated in Equation 1. Lighter colors indicate a larger proportion of the posterior with a reliable non-linear change.

### Model MT_2: Tool-using group: comparing tidal patterns in wet vs. dry seasons

**Table S4.** Posterior mean model estimates of Model MT_2, a Poisson GAM (Bayes R^2^ = 0.03). All estimated population-level effects are on the logit scale. Dry season is the reference category (the intercept).

|  | *Estimate* | *CI_95_low* | *CI_95_high* |
| --- | --- | --- | --- |
| *Smooth terms* |  |  |  |
| Tidetime-Distance_1 | 0.49 | 0.02 | 1.38 |
| Tidetime-Distance_2 | 1.00 | 0.03 | 3.45 |
| Tidetime-Distance_3 | 0.67 | 0.02 | 2.15 |
| Tidetime-Distance_4 | 0.67 | 0.02 | 2.10 |
| Tidetime-Distance dry season_1 | 0.91 | 0.30 | 1.77 |
| Tidetime-Distance dry season_2 | 0.76 | 0.02 | 2.78 |
| Tidetime-Distance dry season_3 | 0.66 | 0.02 | 2.08 |
| Tidetime-Distance wet season_1 | 2.29 | 1.35 | 3.43 |
| Tidetime-Distance wet season_2 | 0.95 | 0.03 | 3.23 |
| Tidetime-Distance wet season_3 | 1.00 | 0.05 | 2.76 |
| Camera location | 0.14 | 0.08 | 0.22 |
| *Population-Level Effects* |  |  |  |
| Intercept | 0.58 | 0.45 | 0.73 |
| Season: Wet | 0.02 | -0.11 | 0.10 |
| Tensor product of Tidetime-Distance | -0.03 | -0.12 | 0.06 |

**Figure S6.**
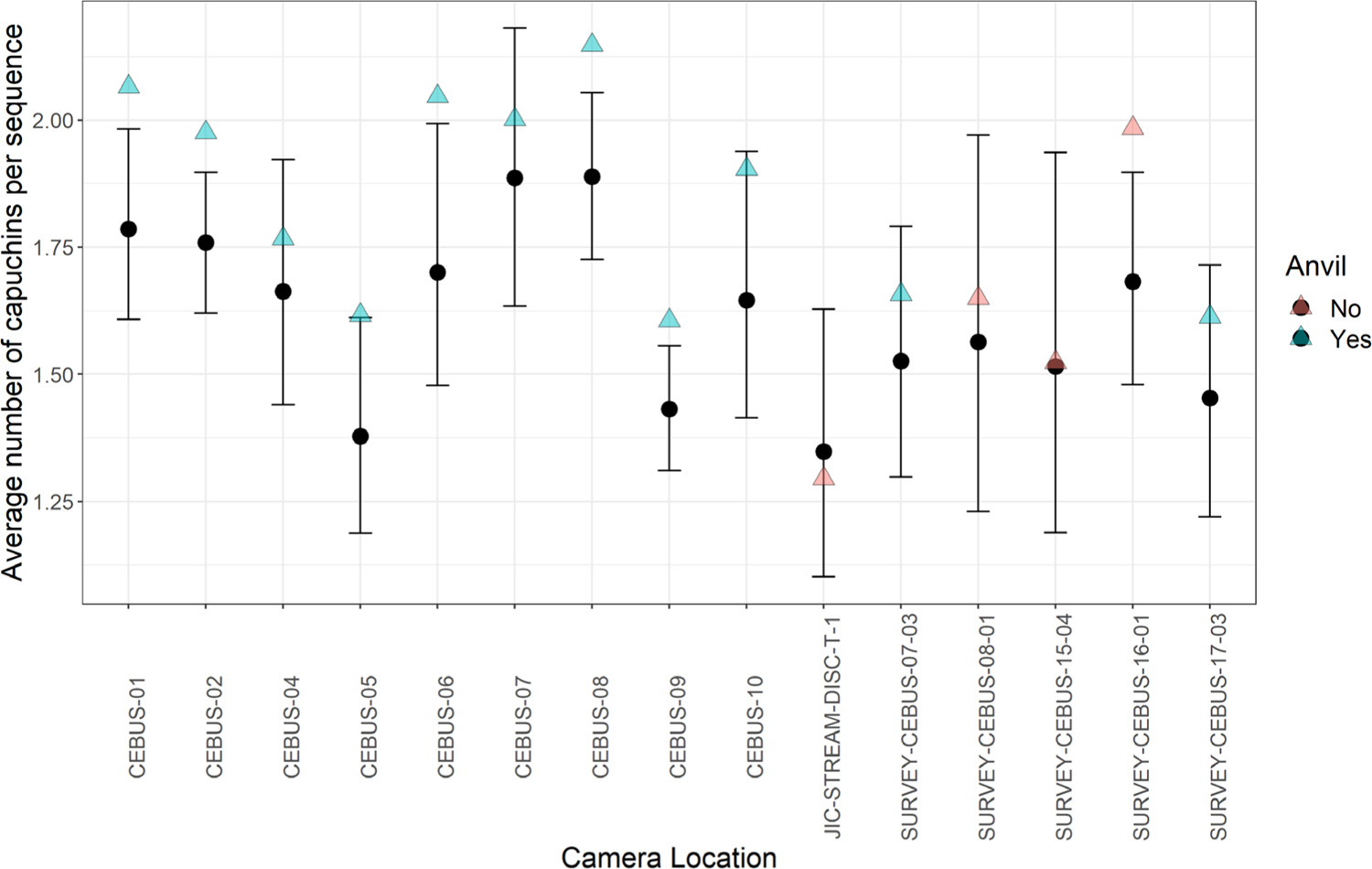
Model MT_2’s posterior mean estimates of the average number of capuchins per camera location represented by black circles. Bars indicate the 95% credible interval. Triangles represent the real average. Blue indicates cameras placed targeted on tool use anvils and red cameras not placed on a targeted tool use location.

**Figure S7.**
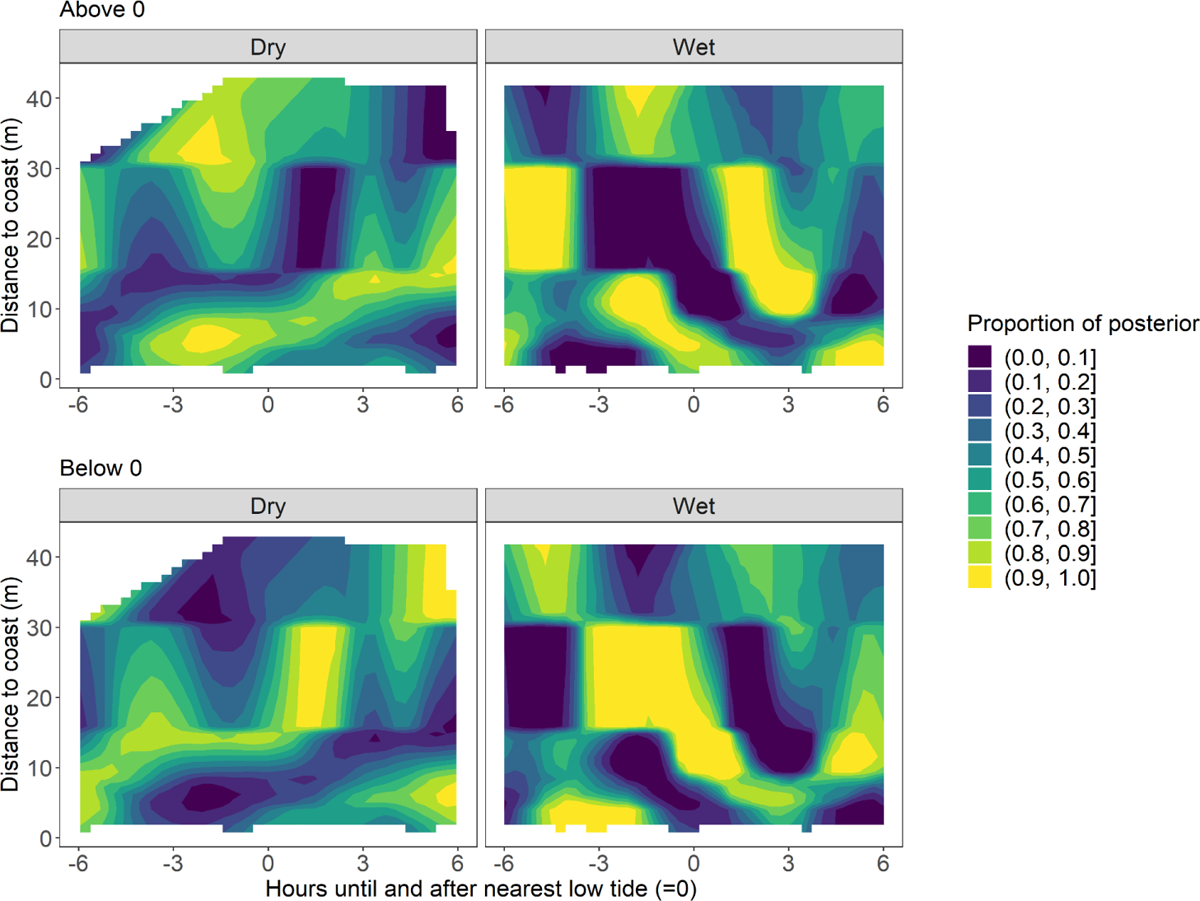
2D heatmap showing for Model MT_2 what proportion of the posterior of the derivative lies above or below 0 as calculated in Equation 1. Lighter colors indicate a larger proportion of the posterior with a reliable non-linear change.

### Model MT_3: Non-tool-using groups: comparing tidal patterns in wet vs. dry season

**Table S5.** Posterior mean model estimates of Model MT_3, a Poisson GAM (Bayes R^2^ = 0.25). All estimated population-level effects are on the logit scale. Dry season is the reference category (the intercept).

|  | *Estimate* | *CI_95_low* | *CI_95_high* |
| --- | --- | --- | --- |
| *Smooth terms* |  |  |  |
| Tidetime-Distance_1 | 0.47 | 0.03 | 1.46 |
| Tidetime-Distance_2 | 0.98 | 0.03 | 3.56 |
| Tidetime-Distance_3 | 0.65 | 0.02 | 2.20 |
| Tidetime-Distance_4 | 0.56 | 0.01 | 1.93 |
| Tidetime-Distance dry season_1 | 4.60 | 2.16 | 6.61 |
| Tidetime-Distance dry season_2 | 0.89 | 0.03 | 3.25 |
| Tidetime-Distance dry season_3 | 1.26 | 0.03 | 7.10 |
| Tidetime-Distance wet season_1 | 0.29 | 0.01 | 0.96 |
| Tidetime-Distance wet season_2 | 0.73 | 0.02 | 2.46 |
| Tidetime-Distance wet season_3 | 0.50 | 0.02 | 1.67 |
| Camera location | 0.28 | 0.18 | 0.43 |
| *Population-Level Effects* |  |  |  |
| Intercept | 0.58 | 0.36 | 0.79 |
| Season: Wet | 0.00 | -0.16 | 0.15 |
| Tensor product of Tidetime-Distance | 0.11 | -0.04 | 0.27 |

**Figure S8.**
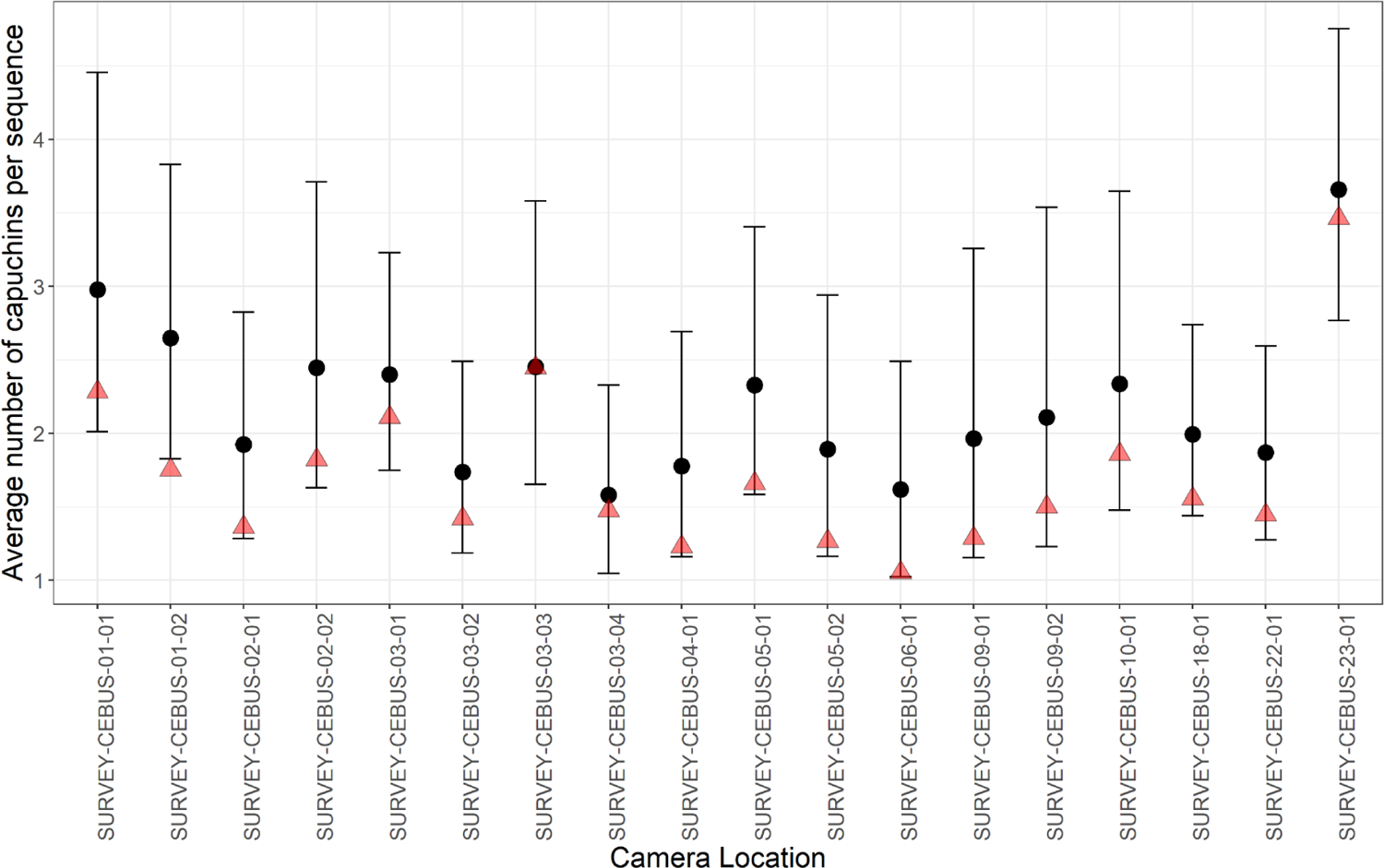
Model MT_3’s posterior mean estimates of the average number of capuchins per camera location represented by black circles. Bars indicate the 95% credible interval. Triangles represent the real average. Blue indicates cameras placed targeted on tool use anvils and red cameras not placed on a targeted tool use location.

**Figure S9.**
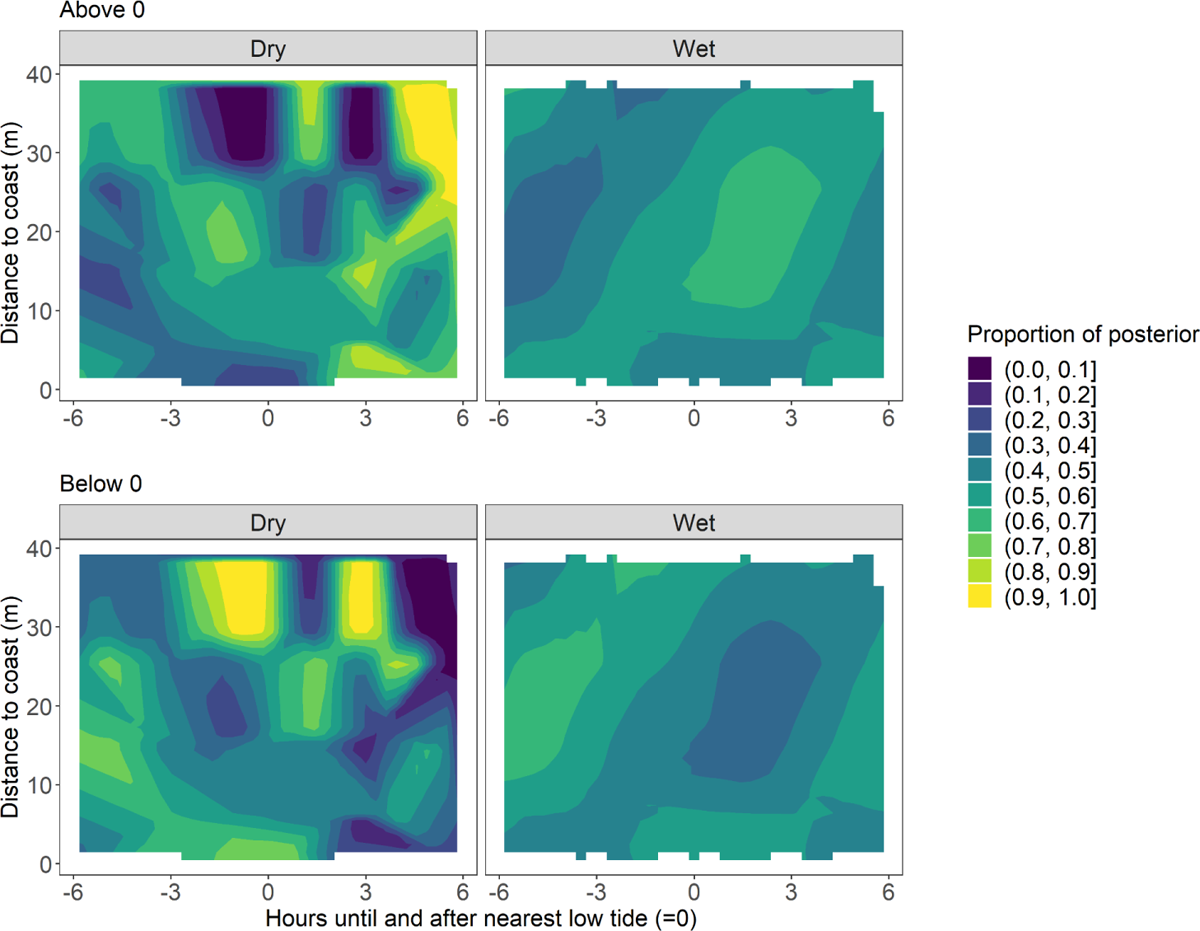
2D heatmap showing for Model MT_3 what proportion of the posterior of the derivative lies above or below 0 as calculated in Equation 1. Lighter colors indicate a larger proportion of the posterior with a reliable non-linear change.

### Model MD_1: Tool-using group: comparing hourly activity in wet vs. dry season

**Table S6.** Posterior mean model estimates of Model MD_1, a Poisson GAM (Bayes R^2^ = 0.04). All estimated population-level effects are on the logit scale. Dry season is the reference category (the intercept).

|  | *Estimate* | *CI_95_low* | *CI_95_high* |
| --- | --- | --- | --- |
| *Smooth terms* |  |  |  |
| Hour-Distance_1 | 0.77 | 0.02 | 2.60 |
| Hour-Distance_2 | 0.97 | 0.03 | 3.46 |
| Hour-Distance_3 | 0.95 | 0.03 | 3.42 |
| Hour-Distance_4 | 0.94 | 0.03 | 3.26 |
| Hour-Distance_5 | 0.91 | 0.02 | 3.30 |
| Hour-Distance dry season_1 | 1.39 | 0.81 | 2.15 |
| Hour-Distance dry season_2 | 0.78 | 0.02 | 2.74 |
| Hour-Distance dry season_3 | 0.73 | 0.02 | 2.31 |
| Hour-Distance wet season_1 | 1.81 | 1.00 | 2.79 |
| Hour-Distance wet season_2 | 0.89 | 0.02 | 3.13 |
| Hour-Distance wet season_3 | 1.18 | 0.04 | 3.37 |
| Camera location | 0.13 | 0.08 | 0.22 |
| *Population-Level Effects* |  |  |  |
| Intercept | 0.60 | 0.43 | 0.78 |
| Season: Wet | -0.04 | -0.20 | 0.13 |
| Tensor product of Hour-Distance_1 | -0.04 | -0.19 | 0.09 |
| Tensor product of Hour-Distance_2 | 0.05 | -0.07 | 0.18 |
| Tensor product of Hour-Distance_3 | 0.02 | -0.09 | 0.13 |

**Figure S10.**
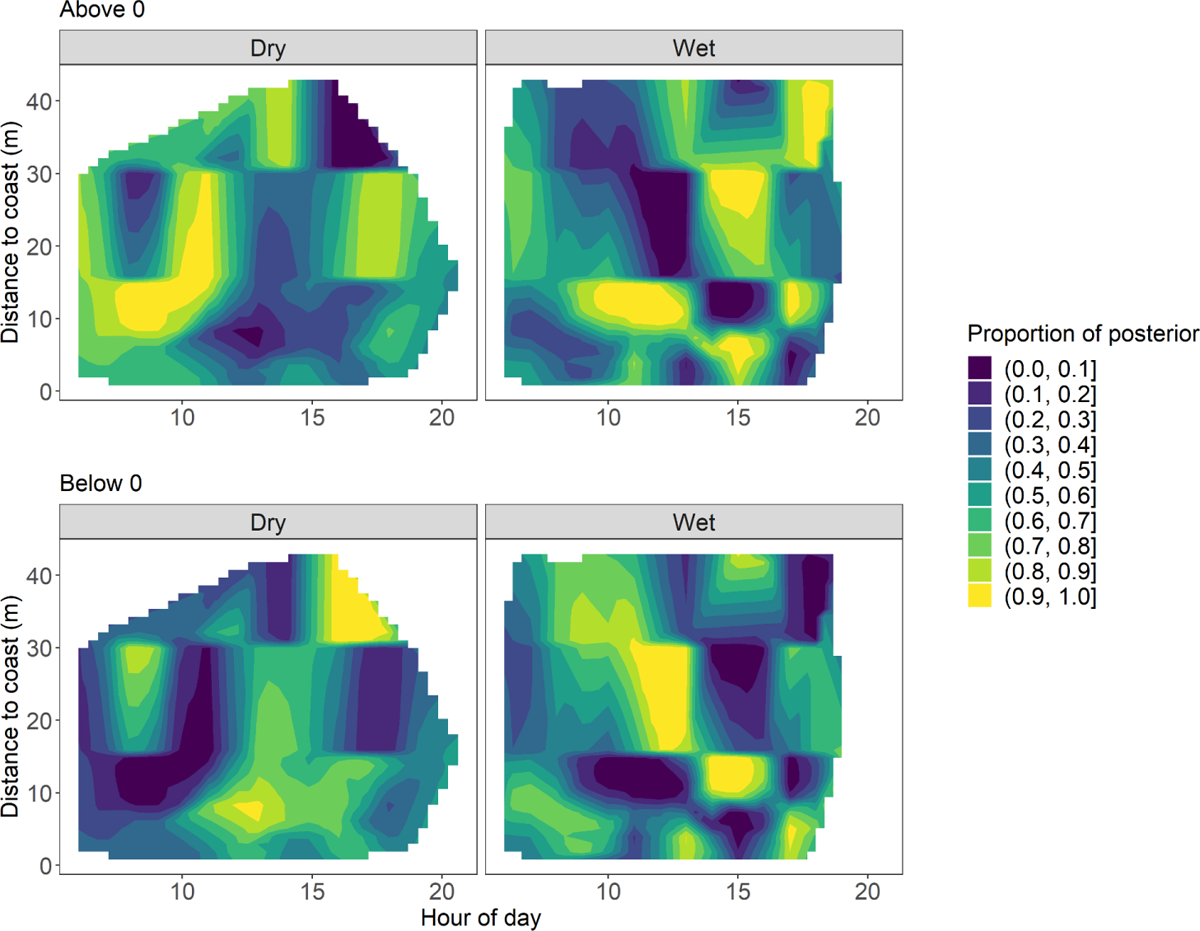
2D heatmap showing for Model MD_1 what proportion of the posterior of the derivative lies above or below 0 as calculated in Equation 1. Lighter colors indicate a larger proportion of the posterior with a reliable non-linear change.

### Model MD_2: Non-tool-using groups: comparing hourly activity in wet vs dry season

**Table S7.**
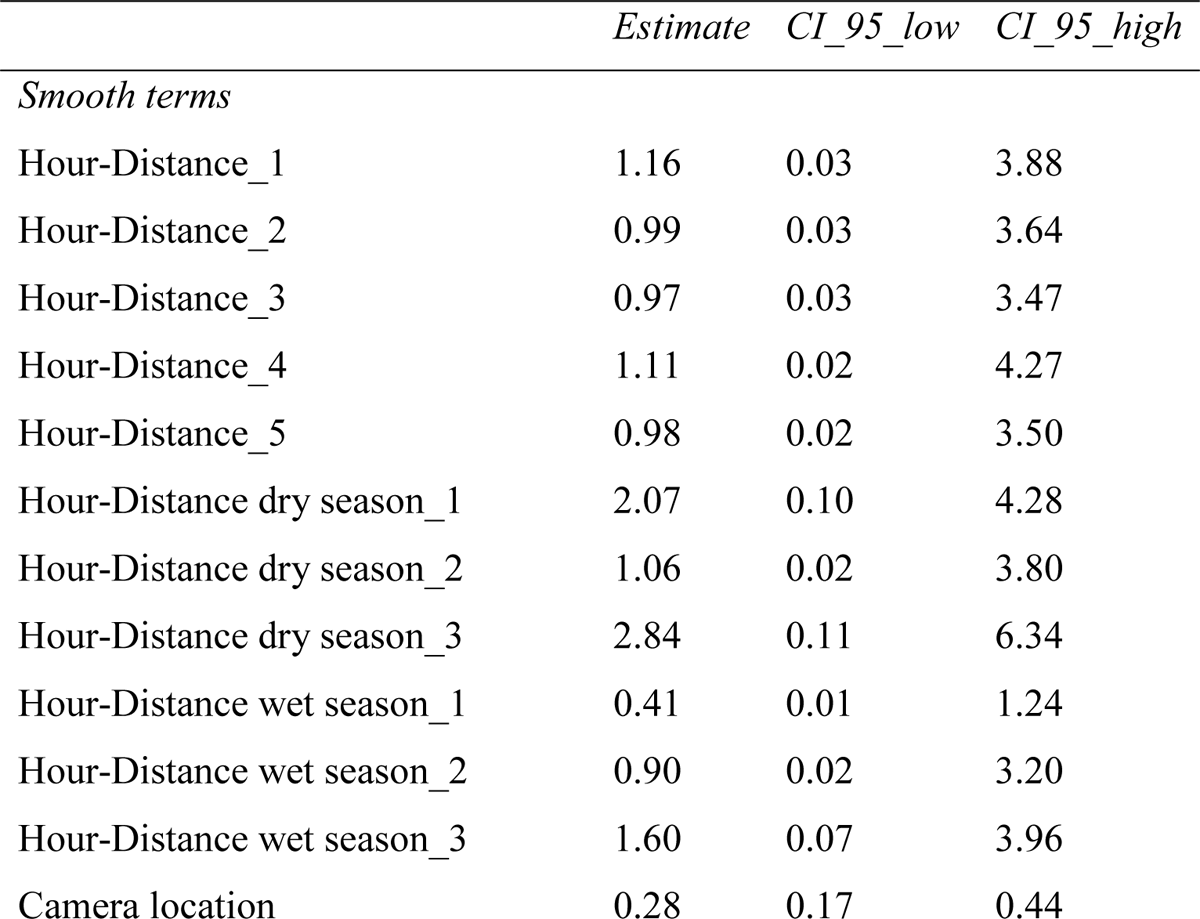
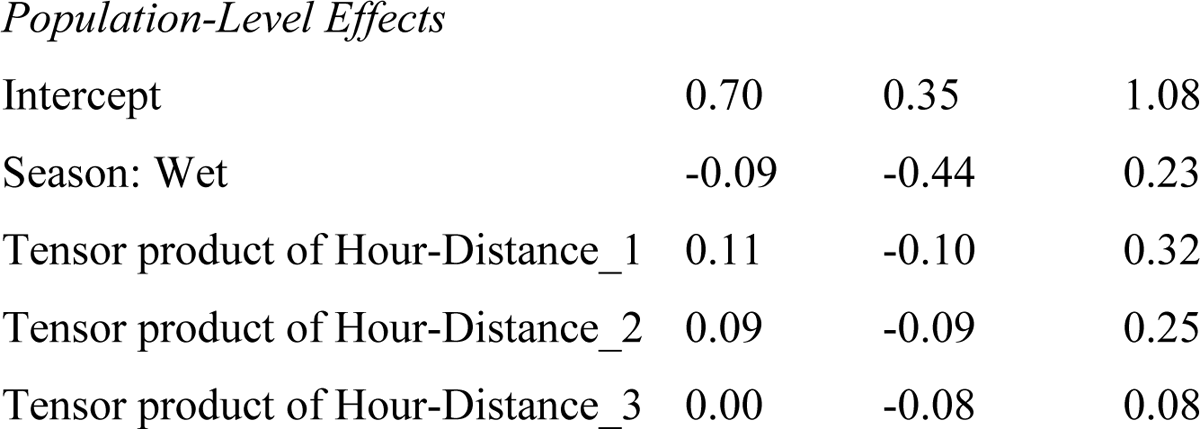
Posterior mean model estimates of Model MD_2, a Poisson GAM (Bayes R^2^ = 0.22). All estimated population-level effects are on the logit scale. Dry season is the reference category (the intercept).

**Model MD 2: Non-tool-using groups: comparing hourly activity in wet vs dry season**
|  | *Estimate* | *CI_95_low* | *CI_95_high* |
| --- | --- | --- | --- |
| *Smooth terms* |  |  |  |
| Hour-Distance_1 | 1.16 | 0.03 | 3.88 |
| Hour-Distance_2 | 0.99 | 0.03 | 3.64 |
| Hour-Distance_3 | 0.97 | 0.03 | 3.47 |
| Hour-Distance_4 | 1.11 | 0.02 | 4.27 |
| Hour-Distance_5 | 0.98 | 0.02 | 3.50 |
| Hour-Distance dry season_1 | 2.07 | 0.10 | 4.28 |
| Hour-Distance dry season_2 | 1.06 | 0.02 | 3.80 |
| Hour-Distance dry season_3 | 2.84 | 0.11 | 6.34 |
| Hour-Distance wet season_1 | 0.41 | 0.01 | 1.24 |
| Hour-Distance wet season_2 | 0.90 | 0.02 | 3.20 |
| Hour-Distance wet season_3 | 1.60 | 0.07 | 3.96 |
| Camera location | 0.28 | 0.17 | 0.44 |
| Intercept | 0.70 | 0.35 | 1.08 |
| Season: Wet | -0.09 | -0.44 | 0.23 |
| Tensor product of Hour-Distance_1 | 0.11 | -0.10 | 0.32 |
| Tensor product of Hour-Distance_2 | 0.09 | -0.09 | 0.25 |
| Tensor product of Hour-Distance_3 | 0.00 | -0.08 | 0.08 |

**Figure S11.**
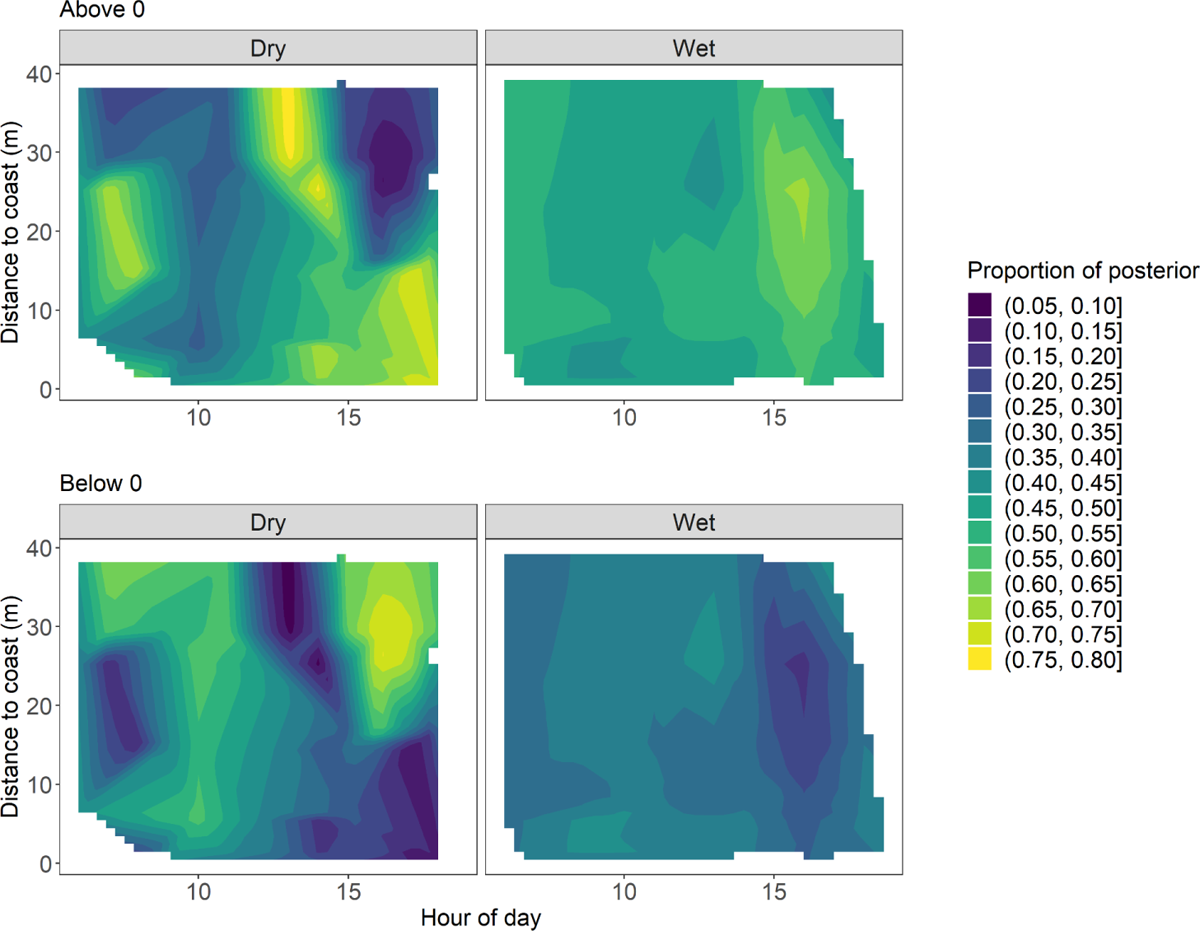
2D heatmap showing for Model MD_2 what proportion of the posterior of the derivative lies above or below 0 as calculated in Equation 1. Lighter colors indicate a larger proportion of the posterior with a reliable non-linear change.

## References

1. Abele, L. G. (1974). Species Diversity of Decapod Crustaceans in Marine Habitats. Ecology, 55(1), 156–161. https://doi.org/10.2307/1934629

2. Alavi, S. (2022). sealavi/GAM-first-derivative-functions: V1.0. Zenodo. https://doi.org/10.5281/zenodo.7457199

3. Araujo, P. E., Carrillo, G. A., Ramirez, W., Blue, S., Chávez, A., Haro, N., García, C., Bravo, N., Castro, S., Hernández, A. H., Prado Wagner, S., & Martin-Solano, S. (2021). First record of tool use by a wild population of Cebus albifrons (Humboldt, 1812) (Primates, Cebidae) in Puerto Misahuallí, Napo, Ecuador. Boletín Técnico, 15(16), Article 16.

4. Barrett, B. J., McElreath, R. L., & Perry, S. E. (2017). Pay-off-biased social learning underlies the diffusion of novel extractive foraging traditions in a wild primate. Proceedings of the Royal Society B: Biological Sciences, 284(1856), 20170358. https://doi.org/10.1098/rspb.2017.0358

5. Barrett, B. J., Monteza-Moreno, C. M., Dogandžić, T., Ibáñez, A., & Crofoot, M. C. (2018). Habitual stone-tool-aided extractive foraging in white-faced capuchins, Cebus capucinus. Royal Society Open Science, 5, 181002. https://doi.org/10.1098/rsos.181002

6. Beck, B. B. (1980). Animal Tool Behaviour: The Use and Manufacture of Tools by Animals. Garland STM Pub.

7. Biro, D., Haslam, M., & Rutz, C. (2013). Tool use as adaptation. Philosophical Transactions of the Royal Society B: Biological Sciences, 368(1630), 20120408. https://doi.org/10.1098/rstb.2012.0408

8. Brakes, P., Carroll, E. L., Dall, S. R. X., Keith, S. A., McGregor, P. K., Mesnick, S. L., Noad, M. J., Rendell, L., Robbins, M. M., Rutz, C., Thornton, A., Whiten, A., Whiting, M. J., Aplin, L. M., Bearhop, S., Ciucci, P., Fishlock, V., Ford, J. K. B., Notarbartolo di Sciara, G., … Garland, E. C. (2021). A deepening understanding of animal culture suggests lessons for conservation. Proceedings. Biological Sciences, 288(1949), 20202718. https://doi.org/10.1098/rspb.2020.2718

9. Bürkner, P.-C. (2017). brms: An R Package for Bayesian Multilevel Models Using Stan. Journal of Statistical Software, 80, 1–28. https://doi.org/10.18637/jss.v080.i01

10. Campbell, J. (2013). White-faced Capuchins (Cebus capucinus) of Cahuita National Park, Costa Rica: Human Foods and Human Interactions. Graduate Theses and Dissertations. https://lib.dr.iastate.edu/etd/13620

11. Campos, F. A., & Fedigan, L. M. (2009). Behavioral adaptations to heat stress and water scarcity in white-faced capuchins (Cebus capucinus) in Santa Rosa National Park, Costa Rica. American Journal of Physical Anthropology, 138(1), 101–111. https://doi.org/10.1002/ajpa.20908

12. Caravaggi, A., Burton, A. C., Clark, D. A., Fisher, J. T., Grass, A., Green, S., Hobaiter, C., Hofmeester, T. R., Kalan, A. K., Rabaiotti, D., & Rivet, D. (2020). A review of factors to consider when using camera traps to study animal behavior to inform wildlife ecology and conservation. Conservation Science and Practice, 2(8), e239. https://doi.org/10.1111/csp2.239

13. Cardiel, J. M., Castroviejo, S., & Velayos, M. (1997). El Parque Nacional de Coiba. El medio físico. In Flora y fauna del Parque Nacional de Coiba (Panamá): Inventario preliminar / editor científico Santiago Castroviejo ; editor adjunto Mauricio Velayos (pp. 11–24). Agencia Española de Cooperación Internacional. https://www.si.edu/object/siris_sil_524758

14. Carlton, J. T., & Hodder, J. (2003). Maritime mammals: Terrestrial mammals as consumers in marine intertidal communities. Marine Ecology Progress Series, 256, 271–286. https://doi.org/10.3354/meps256271

15. Casaer, J., Milotic, T., Liefting, Y., Desmet, P., & Jansen, P. (2019). Agouti: A platform for processing and archiving of camera trap images. Biodiversity Information Science and Standards, 3, e46690. https://doi.org/10.3897/biss.3.46690

16. Clavel, J., Julliard, R., & Devictor, V. (2011). Worldwide decline of specialist species: Toward a global functional homogenization? Frontiers in Ecology and the Environment, 9(4), 222–228. https://doi.org/10.1890/080216

17. Collin, R., Kerr, K., Contolini, G., & Ochoa, I. (2017). Reproductive cycles in tropical intertidal gastropods are timed around tidal amplitude cycles. Ecology and Evolution, 7(15), 5977–5991. https://doi.org/10.1002/ece3.3166

18. Collin, R., & Ochoa, I. (2016). Influence of seasonal environmental variation on the reproduction of four tropical marine gastropods. Marine Ecology Progress Series, 555, 125–139. https://doi.org/10.3354/meps11815

19. Conradt, L. (2000). Use of a seaweed habitat by red deer (Cervus elaphus L.). Journal of Zoology, 250(4), 541–549. https://doi.org/10.1111/j.1469-7998.2000.tb00795.x

20. Crawford, M. A., & Broadhurst, C. L. (2012). The role of docosahexaenoic and the marine food web as determinants of evolution and hominid brain development: The challenge for human sustainability. Nutrition and Health, 21(1), 17–39. https://doi.org/10.1177/0260106012437550

21. Crofoot, M. (2007). Mating and feeding competition in white-faced capuchins (Cebus capucinus): The importance of short- and long-term strategies. Behaviour, 144(12), 1473–1495. https://doi.org/10.1163/156853907782512119

22. Curtis, C. J., & Simpson, G. L. (2014). Trends in bulk deposition of acidity in the UK, 1988– 2007, assessed using additive models. Ecological Indicators, 37, 274–286. https://doi.org/10.1016/j.ecolind.2012.10.023

23. de Chevalier, G., Bouret, S., Bardo, A., Simmen, B., Garcia, C., & Prat, S. (2022). Cost-Benefit Trade-Offs of Aquatic Resource Exploitation in the Context of Hominin Evolution. Frontiers in Ecology and Evolution, 10. https://doi.org/10.3389/fevo.2022.812804

24. De Vynck, J. C., Anderson, R., Atwater, C., Cowling, R. M., Fisher, E. C., Marean, C. W., Walker, R. S., & Hill, K. (2016). Return rates from intertidal foraging from Blombos Cave to Pinnacle Point: Understanding early human economies. Journal of Human Evolution, 92, 101–115. https://doi.org/10.1016/j.jhevol.2016.01.008

25. Dos Santos, R. R., Sousa, A. A. de Fragaszy, D. M., & Ferreira, R. G. (2019). The Role of Tools in the Feeding Ecology of Bearded Capuchins Living in Mangroves. In K. Nowak, A. A. Barnett, & I. Matsuda (Eds.), Primates in Flooded Habitats (1st ed., pp. 59–63). Cambridge University Press. https://doi.org/10.1017/9781316466780.010

26. Eibl-Eibesfeldt, I. (1961). Über den Werkzeuggebrauch des Spechtfinken Camarhynchus pallidus. Zeitschrift Für Tierpsychologie, 18(3), 343–346. https://doi.org/10.1111/j.1439-0310.1961.tb00424.x

27. Erlandson, J. M. (2001). The Archaeology of Aquatic Adaptations: Paradigms for a New Millennium. Journal of Archaeological Research, 9(4), 287–350. https://doi.org/10.1023/A:1013062712695

28. Fedigan, L. M., & Jack, K. M. (2012). Tracking Neotropical Monkeys in Santa Rosa: Lessons from a Regenerating Costa Rican Dry Forest. In P. M. Kappeler & D. P. Watts (Eds.), Long-Term Field Studies of Primates (pp. 165–184). Springer. https://doi.org/10.1007/978-3-642-22514-7_8

29. Garrity, S. D. (1984). Some Adaptations of Gastropods to Physical Stress on a Tropical Rocky Shore. Ecology, 65(2), 559–574. https://doi.org/10.2307/1941418

30. Gaylord, B., Kroeker, K. J., Sunday, J. M., Anderson, K. M., Barry, J. P., Brown, N. E., Connell, S. D., Dupont, S., Fabricius, K. E., Hall-Spencer, J. M., Klinger, T., Milazzo, M., Munday, P. L., Russell, B. D., Sanford, E., Schreiber, S. J., Thiyagarajan, V., Vaughan, M. L. H., Widdicombe, S., & Harley, C. D. G. (2015). Ocean acidification through the lens of ecological theory. Ecology, 96(1), 3–15. https://doi.org/10.1890/14-0802.1

31. Goodall, J. (1964). Tool-Using and Aimed Throwing in a Community of Free-Living Chimpanzees. Nature, 201(4926), Article 4926. https://doi.org/10.1038/2011264a0

32. Goodman, M., Hayward, T., & Hunt, G. R. (2018). Habitual tool use innovated by free-living New Zealand kea. Scientific Reports, 8(1), Article 1. https://doi.org/10.1038/s41598-018-32363-9

33. Gumert, M. D., Kluck, M., & Malaivijitnond, S. (2009). The physical characteristics and usage patterns of stone axe and pounding hammers used by long-tailed macaques in the Andaman Sea region of Thailand. American Journal of Primatology, 71(7), 594– 608. https://doi.org/10.1002/ajp.20694

34. Gumert, M. D., & Malaivijitnond, S. (2012). Marine prey processed with stone tools by burmese long-tailed macaques (Macaca fascicularis aurea) in intertidal habitats. American Journal of Physical Anthropology, 149(3), 447–457. https://doi.org/10.1002/ajpa.22143

35. Gyory, J. (2010). Larval ecology and synchronous reproduction of two crustacean species: Semibalanus balanoides in New England, USA and Gecarcinus quadratus in Veraguas, Panama [Thesis, Massachusetts Institute of Technology]. https://dspace.mit.edu/handle/1721.1/62786

36. Henke-von der Malsburg, J., Kappeler, P. M., & Fichtel, C. (2020). Linking ecology and cognition: Does ecological specialisation predict cognitive test performance? Behavioral Ecology and Sociobiology, 74(12), 154. https://doi.org/10.1007/s00265-020-02923-z

37. Henke-von der Malsburg, J., Kappeler, P. M., & Fichtel, C. (2021). Linking cognition to ecology in wild sympatric mouse lemur species. Proceedings of the Royal Society B: Biological Sciences, 288(1963), 20211728. https://doi.org/10.1098/rspb.2021.1728

38. Hijmans, R. J. (2022). terra: Spatial Data Analysis (1.5-21) [R Package]. https://CRAN.R-project.org/package=terra

39. Hill, S. E., & Winder, I. C. (2019). Predicting the impacts of climate change on Papio baboon biogeography: Are widespread, generalist primates ‘safe’? Journal of Biogeography, 46(7), 1380–1405. https://doi.org/10.1111/jbi.13582

40. Ibáñez, A. (2011). Guía botánica del Parque nacional Coiba. International Cooperative Biodiversity Groups: Instituo Smithsonian de Investigaciones Tropicals: Secretaría Nacional de C.

41. Ibáñez, A., Pérez-Jordá, J., Juste, J., & Guillén, A. (1997). Los mamíferos terrestres del Parque Nacional Coiba (Panamá) (ed. S Castroviejo, pp. 469–486). Agencia Española de Cooperacíon Internacional.

42. IPCC. (2022). Climate Change 2022: Impacts, Adaptation, and Vulnerability. Cambridge University Press.

43. Irons, D. B. (1998). Foraging Area Fidelity of Individual Seabirds in Relation to Tidal Cycles and Flock Feeding. Ecology, 79(2), 647–655. https://doi.org/10.1890/0012-9658(1998)079[0647:FAFOIS]2.0.CO;2

44. Isaza, I., & Vrba, E. (2010). Ocupación pre-colombina de las Islas del Parque Nacional Coiba. Panama: Informe final del Proyecto PRB08-003 archivos SENACYT.

45. Izar, P., Peternelli-dos-Santos, L., Rothman, J. M., Raubenheimer, D., Presotto, A., Gort, G., Visalberghi, E. M., & Fragaszy, D. M. (2022). Stone tools improve diet quality in wild monkeys. Current Biology, 32(18), 4088–4092.e3. https://doi.org/10.1016/j.cub.2022.07.056

46. Joordens, J. C. A., Kuipers, R. S., Wanink, J. H., & Muskiet, F. A. J. (2014). A fish is not a fish: Patterns in fatty acid composition of aquatic food may have had implications for hominin evolution. Journal of Human Evolution, 77, 107–116. https://doi.org/10.1016/j.jhevol.2014.04.004

47. Krützen, M., Kreicker, S., MacLeod, C. D., Learmonth, J., Kopps, A. M., Walsham, P., & Allen, S. J. (2014). Cultural transmission of tool use by Indo-Pacific bottlenose dolphins (Tursiops sp.) provides access to a novel foraging niche. Proceedings of the Royal Society B: Biological Sciences, 281(1784), 20140374. https://doi.org/10.1098/rspb.2014.0374

48. Krutzen, M., Mann, J., Heithaus, M. R., Connor, R. C., Bejder, L., & Sherwin, W. B. (2005). Cultural transmission of tool use in bottlenose dolphins. Proceedings of the National Academy of Sciences, 102(25), 8939–8943. https://doi.org/10.1073/pnas.0500232102

49. Kuwagata, T., Kondo, J., & Sumioka, M. (1994). Thermal effect of the sea breeze on the structure of the boundary layer and the heat budget over land. Boundary-Layer Meteorology, 67(1), 119–144. https://doi.org/10.1007/BF00705510

50. Levings, S. C., & Garrity, S. D. (1983). Diel and tidal movement of two co-occurring neritid snails; differences in grazing patterns on a tropical rocky shore. Journal of Experimental Marine Biology and Ecology, 67(3), 261–278. https://doi.org/10.1016/0022-0981(83)90043-6

51. Lewis, M. C., & O’Riain, M. J. (2017). Foraging Profile, Activity Budget and Spatial Ecology of Exclusively Natural-Foraging Chacma Baboons (Papio ursinus) on the Cape Peninsula, South Africa. International Journal of Primatology, 38(4), 751–779. https://doi.org/10.1007/s10764-017-9978-5

52. Lewis, M. C., & Sealy, J. C. (2018). Coastal complexity: Ancient human diets inferred from Bayesian stable isotope mixing models and a primate analogue. PLOS ONE, 13(12), e0209411. https://doi.org/10.1371/journal.pone.0209411

53. Mannino, M. A., & Thomas, K. D. (2002). Depletion of a resource? The impact of prehistoric human foraging on intertidal mollusc communities and its significance for human settlement, mobility and dispersal. World Archaeology, 33(3), 452–474. https://doi.org/10.1080/00438240120107477

54. Marean, C. W. (2014). The origins and significance of coastal resource use in Africa and Western Eurasia. Journal of Human Evolution, 77, 17–40. https://doi.org/10.1016/j.jhevol.2014.02.025

55. Marean, C. W., Bar-Matthews, M., Bernatchez, J., Fisher, E., Goldberg, P., Herries, A. I. R., Jacobs, Z., Jerardino, A., Karkanas, P., Minichillo, T., Nilssen, P. J., Thompson, E., Watts, I., & Williams, H. M. (2007). Early human use of marine resources and pigment in South Africa during the Middle Pleistocene. Nature, 449(7164), 905–908. https://doi.org/10.1038/nature06204

56. Marshall, A. J., Boyko, C. M., Feilen, K. L., Boyko, R. H., & Leighton, M. (2009). Defining fallback foods and assessing their importance in primate ecology and evolution. American Journal of Physical Anthropology, 140(4), 603–614. https://doi.org/10.1002/ajpa.21082

57. Marshall, A. J., & Wrangham, R. W. (2007). Evolutionary Consequences of Fallback Foods. International Journal of Primatology, 28(6), 1219–1235. https://doi.org/10.1007/s10764-007-9218-5

58. Mazzia, N., & Flegenheimer, N. (2015). Detailed fatty acids analysis on lithic tools, Cerro El Sombrero Cima, Argentina. Quaternary International, 363, 94–106. https://doi.org/10.1016/j.quaint.2014.04.027

59. Méndez-Carvajal, P. G., & Valdés-Díaz, S. (2017). Use of anvils and other feeding behaviour observed in cebus imitator, Coiba island, Panama. Tecnociencia, 19(1), 5–18.

60. Milton, K., & Mittermeier, R. A. (1977). A brief survey of the primates of Coiba Island, Panama. Primates, 18(4), 931–936. https://doi.org/10.1007/BF02382942

61. Monteith, D. T., Evans, C. D., Henrys, P. A., Simpson, G. L., & Malcolm, I. A. (2014). Trends in the hydrochemistry of acid-sensitive surface waters in the UK 1988–2008. Ecological Indicators, 37, 287–303. https://doi.org/10.1016/j.ecolind.2012.08.013

62. Monteza-Moreno, C. M., Crofoot, M. C., Grote, M. N., & Jansen, P. A. (2020). Increased terrestriality in a Neotropical primate living on islands with reduced predation risk. Journal of Human Evolution, 143, 102768. https://doi.org/10.1016/j.jhevol.2020.102768

63. Monteza-Moreno, C. M., Dogandžić, T., McLean, K. A., Castillo-Caballero, P. L., Mijango-Ramos, Z., Del Rosario-Vargas, E., Crofoot, M. C., & Barrett, B. J. (2020). White-Faced Capuchin, Cebus capucinus imitator, Hammerstone and Anvil Tool Use in Riparian Habitats on Coiba Island, Panama. International Journal of Primatology, 41(3), 429–433. https://doi.org/10.1007/s10764-020-00156-5

64. Moynihan, M. (1976). The New World Primates: Adaptive Radiation and the Evolution of Social Behavior, Languages, and Intelligence. Princeton University Press. https://www.jstor.org/stable/j.ctt13x16qs

65. Navarrete, S., & Castilla, J. (1993). Predation by Norway rats in the intertidal zone of central Chile. Marine Ecology Progress Series, 92, 187–199. https://doi.org/10.3354/meps092187

66. Nielsen, S. M. (1991). Fishing Arctic foxes Alopex lagopus on a rocky island in west Greenland. Polar Research, 9(2), Article 2. https://doi.org/10.3402/polar.v9i2.6793

67. Oppenheimer, J. R. (1968). Behavior and ecology of the white-faced capuchin monkey, Cebus capucinus, on Barro Colorado Island, Canal Zone. University of Illinois at Urbana-Champaign.

68. Panger, M. A., Perry, S., Rose, L., Gros-Louis, J., Vogel, E., Mackinnon, K. C., & Baker, M. (2002). Cross-site differences in foraging behavior of white-faced capuchins (Cebus capucinus). American Journal of Physical Anthropology, 119(1), 52–66. https://doi.org/10.1002/ajpa.10103

69. Pedersen, E. J., Miller, D. L., Simpson, G. L., & Ross, N. (2019). Hierarchical generalized additive models in ecology: An introduction with mgcv. PeerJ, 7, e6876. https://doi.org/10.7717/peerj.6876

70. Perez, R., & Condit, R. (n.d.). Tree Atlas of Panama. Center for Tropical Forest Science, Smithsonian Tropical Research Institute. Accessed December 2022.

71. Pérez, R., Condit, R., Aguilar, S., Hernández, A., & Villareal, A. (1996). Inventario de la vegetación de la isla de Coiba, Panamá: Composición y florística. Revista de Biologia Tropical, 44(1), Article 1.

72. Perry, S. (2011). Social traditions and social learning in capuchin monkeys (Cebus). Philosophical Transactions of the Royal Society B: Biological Sciences, 366(1567), 988–996. https://doi.org/10.1098/rstb.2010.0317

73. Perry, S., Godoy, I., & Lammers, W. (2012). The Lomas Barbudal Monkey Project: Two Decades of Research on Cebus capucinus. In P. M. Kappeler & D. P. Watts (Eds.), Long-Term Field Studies of Primates (pp. 141–163). Springer. https://doi.org/10.1007/978-3-642-22514-7_7

74. Perry, S., & Ordonez Jiménez, J. (2006). The effects of food size, rarity, and processing complexity on white-faced capuchins visual attention to foraging conspecifics. In G. Hohmann, M. Robbins, & C. Boesch (Eds.), Feeding Ecology in Apes and other Primates (pp. 203–234). Cambridge University Press. https://hdl.handle.net/11858/00-001M-0000-0010-00A2-6

75. Planet Team. (2022). Planet Application Program Interface: In Space for Life on Earth. https://api.planet.com.

76. R Core Team. (2022). R: A language and environment for statistical computing. R Foundation for Statistical Computing. https://www.R-project.org/

77. Ridout, M. S., & Linkie, M. (2009). Estimating overlap of daily activity patterns from camera trap data. Journal of Agricultural, Biological, and Environmental Statistics, 14(3), 322–337. https://doi.org/10.1198/jabes.2009.08038

78. Rowcliffe, J. M., Kays, R., Kranstauber, B., Carbone, C., & Jansen, P. A. (2014). Quantifying levels of animal activity using camera trap data. Methods in Ecology and Evolution, 5(11), 1170–1179. https://doi.org/10.1111/2041-210X.12278

79. Rutz, C., Klump, B. C., Komarczyk, L., Leighton, R., Kramer, J., Wischnewski, S., Sugasawa, S., Morrissey, M. B., James, R., St Clair, J. J. H., Switzer, R. A., & Masuda, B. M. (2016). Discovery of species-wide tool use in the Hawaiian crow. Nature, 537(7620), Article 7620. https://doi.org/10.1038/nature19103

80. Rutz, C., & St Clair, J. J. H. (2012). The evolutionary origins and ecological context of tool use in New Caledonian crows. Behavioural Processes, 89(2), 153–165. https://doi.org/10.1016/j.beproc.2011.11.005

81. Sanz, C. M., & Morgan, D. B. (2013). Ecological and social correlates of chimpanzee tool use. Philosophical Transactions of the Royal Society B: Biological Sciences, 368(1630), 20120416. https://doi.org/10.1098/rstb.2012.0416

82. Shumaker, R. W., Walkup, K. R., Beck, B. B., & Burghardt, G. M. (2011). Animal Tool Behavior: The Use and Manufacture of Tools by Animals (revised and updated edition). Johns Hopkins University Press.

83. Simmons, B. L., Sterling, J., & Watson, J. C. (2014). Species and size-selective predation by raccoons (Procyon lotor) preying on introduced intertidal clams. Canadian Journal of Zoology, 92(12), 1059–1065. https://doi.org/10.1139/cjz-2014-0108

84. Sollmann, R. (2018). A gentle introduction to camera-trap data analysis. African Journal of Ecology, 56(4), 740–749. https://doi.org/10.1111/aje.12557

85. Stander, P. E. (2019). Lions (Panthera leo) specialising on a marine diet in the Skeleton Coast Park, Namibia. Namibian Journal of Environment, 3, A-10.

86. Steward, J. (2006). The concept and method of cultural ecology. In N. Haenn & R. Wilk (Eds.), The Environment in Anthropology: A Reader in Ecology, Culture, and Sustainable Living (pp. 5–9). NYU Press.

87. Suraci, J. P., Clinchy, M., & Zanette, L. Y. (2017). Do Large Carnivores and Mesocarnivores Have Redundant Impacts on Intertidal Prey? PLOS ONE, 12(1), e0170255. https://doi.org/10.1371/journal.pone.0170255

88. Titcomb, M., & O’Dea, A. (2020). Post-glacial Sea Level rise on the Isthmus of Panama. https://doi.org/10.25573/data.11919276.

89. Trowbridge, C. D. (1994). Life at the edge: Population dynamics and salinity tolerance of a high intertidal, pool-dwelling ascoglossan opisthobranch on New Zealand rocky shores. Journal of Experimental Marine Biology and Ecology, 182(1), 65–84. https://doi.org/10.1016/0022-0981(94)90211-9

90. Tsuji, Y., & Kazahari, N. (2019). Maritime macaques: Ecological background of seafood eating by wild Japanese macaques (Macaca fuscata). In A. Barnett, I. Matsuda, & K. Nowak (Eds.), Primates in Flooded Habitats: Ecology and Conservation (pp. 135– 143). Cambridge University Press.

91. Wickham, H. (2016). ggplot2: Elegant Graphics for Data Analysis. Springer-Verlag New York. https://ggplot2.tidyverse.org

92. Wood, S. A. (2006). Generalized Additive Models: An Introduction with R. Chapman and Hall/CRC. https://doi.org/10.1201/9781420010404

